# A Kernel-Spectral Index for Single-Neuron Input–Output Complexity

**DOI:** 10.64898/2026.11.04.764153

**Authors:** Zixuan Huang, Wenjie Chen, Bo Lei, Guozhang Chen

## Abstract

The computational capacity of individual neurons in the brain is a central problem in computational neuroscience and an increasingly relevant link between neuroscience and AI. Recent work fitting artificial neural networks to detailed cortical pyramidal neurons showed that reproducing their millisecond- precision input–output behavior can require temporal convolutional networks with five to eight layers, suggesting that cortical neurons may function as deep computational units inside brain circuits. Yet current quantitative studies of single-neuron complexity are largely surrogate-based; consequently, the obtained results are difficult to interpret, and the process often demands massive computational resources. We introduce the Spectral Complexity Index (SCI), a white-box measure that represents presynaptic histories with an explicit kernel geometry and quantifies how many ordered spectral modes are needed to explain somatic response energy. Across 35 biophysically detailed human and mouse cortical neuron models, SCI yields a statistically significant separation of the two species, strongly agrees with the DNN-based Functional Complexity Index on these cells (Spearman correlation: *ρ* = 0.797), and remains stable under sampling and kernel perturbations. Additionally, compared to existing methods such as FCI, the computational cost of SCI is reduced to approximately 1*/*10 or less. SCI turns single-neuron complexity from a black-box surrogate property into an interpretable spectral object, enabling reproducible, fast, and efficient comparisons of neuronal computation across cells and species.

## 1 Introduction

Single neurons in the brain transform distributed synaptic histories into somatic voltages and spikes through dendritic filtering, active conductances, local regenerative events, and voltage-dependent synaptic currents. In cortical pyramidal neurons, dendrites can act as structured nonlinear subunits that support branch-level integration, temporal-order sensitivity, and NMDA-mediated supralinear responses [1–5]. Human cortical neurons add a comparative motivation: they differ from rodent neurons in morphology, compartmentalization, membrane properties, and synaptic physiology [6–9]. A useful scalar complexity measure should therefore compare cells and species while remaining tied to the presynaptic histories that actually drive the neuron.

Recent surrogate-based approaches have made this problem measurable. Deep temporal convolutional networks can approximate detailed pyramidal-cell input–output (I/O) maps, and the Functional Complexity Index (FCI) converts the prediction difficulty of spikes and soma voltage into a scalar score by fixing a TCN benchmark [10, 11]. Although highly informative, these methods still suffer from a lack of interpretability. Moreover, the extensive computational overhead required by current techniques—often amounting to several single-GPU years—presents tangible barriers to their practical deployment.

**Table 1:** Comparison of different methods. FCI, Functional Complexity Index. TCN, Temporal Convolutional Network. DBNN, Dendritic Bilinear Neural Network. SCI, Spectral Complexity Index.

| Method | Complexity object | Computational cost | Interpretability |
|---|---|---|---|
| DNN [10] | Minimal TCN size | High | Black-box |
| FCI [11] | Fixed-TCN prediction error | High | Black-box |
| DBNN [12] | Fitted response surrogate | Medium | Partially interpretable |
| SCI (**ours**) | Spectral energy cutoff | Low | White-box |

We introduce the Spectral Complexity Index (SCI), a white-box spectral measure of single-neuron I/O complexity under a specified stimulation ensemble. SCI represents presynaptic histories with channels for synapse type, dendritic domain, electrotonic distance, and time lag. A time-decayed Gaussian kernel is defined on those histories, and the centered somatic response function is decomposed in the eigensystem of the induced kernel operator. The index SCI_*θ*_ is the minimum number of leading modes required to explain a fraction *θ* of response energy. Thus, existing methods can be viewed as different probes of how compactly the same underlying I/O map can be represented, while SCI makes the common target explicit as a response-energy spectrum over ordered input-history modes.

SCI is also computationally advantageous. FCI-style benchmarks require large supervised corpora and multi-seed neural-network training; the FCI protocol, for example, uses 1.2 ×10^4^ ten-second simulations per model [11]. In contrast, only 10^2^–10^3^ ten-second simulations are needed to calculate SCI, and once simulated traces are available, computing five random-seed SCI estimates for one cell takes roughly three minutes in our implementation.

Our contributions are threefold. First, we formulate SCI as a white-box, interpretable complexity index for single-neuron I/O maps, replacing an implicit learned benchmark with visible kernel modes, coefficients, and energy cutoffs. Second, SCI requires substantially fewer computational resources, which provides a fast and efficient method for evaluating neuronal I/O complexity. Third, we evaluate SCI on 35 human and mouse cortical neuron models, showing statistically significant species separation, strong agreement with FCI, robustness to sampling and kernel perturbations, and meaningful association with dendritic morphology.

## 2 Related work

### Dendritic computation and surrogate benchmarks

Dendritic computation provides the biological basis for single-neuron complexity: active dendrites, nonlinear subunits, and local synaptic mechanisms can make the somatic response depend on fine spatial and temporal input structure [1, 3–5]. Neural-network surrogates turn this intuition into measurable prediction problems. Beniaguev et al. showed that detailed pyramidal-cell dynamics can require deep temporal convolutional networks, and Aizenbud et al. defined FCI using a fixed TCN benchmark to compare cortical neurons [10, 11]. SCI is complementary: it does not replace these findings, but gives their notion of approximation difficulty a white-box spectral coordinate system.

### Structured models and kernel spectra

Hierarchical linear–nonlinear models, dendritic bilinear neural networks, and reduced compartmental models improve interpretability by imposing structure on the response model [12–14]. SCI addresses the adjacent question of measurement rather than prediction: it quantifies the number of explicitly ordered input-history modes required to account for response energy. Kernel eigensystems provide the needed coordinates by converting a similarity function into an ordered basis under the input distribution [15, 16]. In SCI, this makes the assumptions of the complexity measure inspectable rather than embedded in a trained surrogate. Simultaneously, SCI enables a further substantial reduction in the required computational resources.

**Figure 1.**
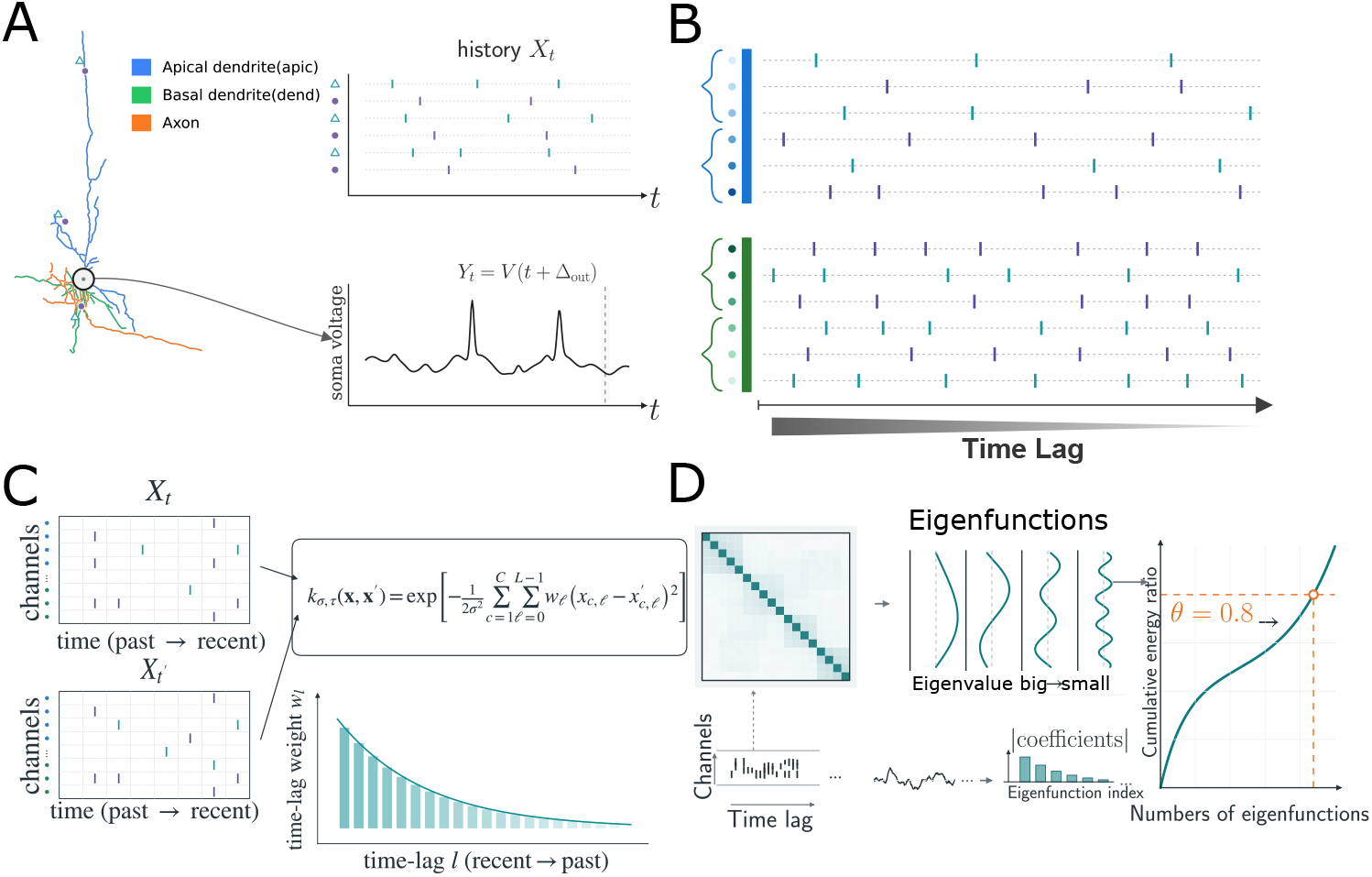
Overview of the Spectral Complexity Index (SCI). **A**, Distributed synaptic input to a compartmental neuron defines a finite-history input variable *X*_*t*_, paired with the next-bin somatic voltage response *Y*_*t*_ = *V* (*t* + Δ_out_). **B**, Presynaptic histories are represented in a shared channel-by-lag coordinate system across morphologies. **C**, A time-weighted Gaussian kernel compares input histories, with stronger weighting on recent lags. **D**, The empirical kernel eigenspectrum provides spectral coordinates for the centered voltage response; SCI_*θ*_ is the minimum number of leading eigenfunctions required to reach spectral-energy cutoff *θ*.

## 3 Spectral complexity index

SCI measures the complexity of a neuron’s input–output map relative to an explicit geometry on presynaptic histories. For a neuron *m*, let *X*_*t*_ ∈*X* ⊆ ℝ^*C×L*^ be the finite presynaptic-history random variable preceding time *t*, and let **x** denote one realization. Here *C* is the number of input channels, and *L* is the length of history. Let *Y*_*t*_ denote a scalar somatic response. In the main analysis, *Y*_*t*_ = *V* (*t* + Δ_out_), the next-bin somatic membrane voltage, i.e. Δ_out_ = *δt* and *δt* = 1 ms is the input temporal resolution. In the experiments below, the firing rates of different cells are all calibrated to [0.8, 1.2] Hz. SCI is defined for the finite-history response map induced by the chosen input representation. Throughout the empirical analysis, we use the working approximation *Y*_*t*_ ≈ *f*_*m*_(*X*_*t*_), so residual variation from earlier unobserved history and latent biophysical state is assumed small for the chosen voltage readout and history window. We assume *f*_*m*_ ∈ *L*^2^(*ρ*).

### 3.1 Presynaptic histories

Each history is represented by channelized excitatory and inhibitory input events. A channel is indexed by

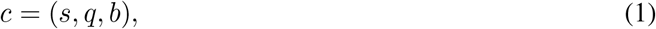

where *s* ∈ {E, I}is the synapse type, ∈ {*q}*basal, apical is the dendritic domain, and *b* ∈ {1, …, 50}is a bin of normalized electrotonic distance from the soma. Here, the distance is normalized per cell by setting the maximum electrotonic distance among all its synapses to 1. Thus *C* = 2× 2× 50 = 200. The coordinate *x*_*c,l*_ records the input event count in channel *c* at lag *l*, with *l* = 0 corresponding to the most recent input bin. This representation keeps the input variable external to the neuron: synaptic conductances, dendritic morphology, and membrane nonlinearities affect SCI through the response *Y*_*t*_, not through a learned hidden state.

For a fixed stimulation protocol, the random histories *X*_*t*_ induce a distribution *ρ* on *X*. The neuronal response is represented by the finite-history conditional response function

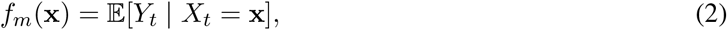

and SCI is defined for *f*_*m*_, *ρ*, and the kernel specified below. In the empirical estimator, the observed voltage sample *y*_*i*_ is treated as a finite-sample proxy for *f*_*m*_(*x*_*i*_) under the working approximation *Y*_*t*_ ≈ *f*_*m*_(*X*_*t*_). Across cells, the stimulation rule is standardized, but the induced distribution can be cell-specific because input calibration and morphology are cell-specific; when necessary, *ρ* should be read as *ρ*_*m*_, with the subscript suppressed for readability.

### 3.2 A time-weighted kernel on input histories

SCI uses a time-decayed Gaussian kernel on the flattened history vector **x** = (*x*_*c,l*_)_*c,l*_ ∈ ℝ^*CL*^. The kernel compares two histories by a weighted squared distance over input channel *c* and lag *l*:

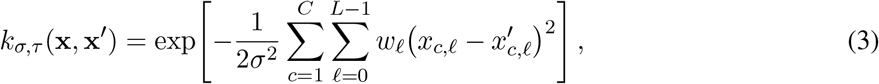

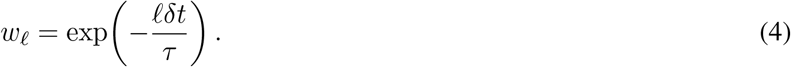

The scalar *σ >* 0 sets the overall bandwidth of the kernel in the channelized input space. The scalar *τ >* 0 sets the decay scale of the temporal metric, and *w*_*l*_ is the dimensionless weight assigned to lag *l*, with *l* = 0 denoting the most recent input bin. See Appendix C.6 for evidence supporting the weighted Gaussian kernel over a linear kernel. Equivalently, the coordinate *x*_*c,l*_ is compared with an effective length scale

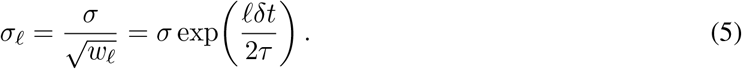

Recent input bins are therefore resolved more finely than older input bins. In all main experiments, we set *σ* = 1 and *τ* = 20 ms, and robustness to these choices is evaluated in Section 5.3 and Appendix C.7. A heuristic explanation for our choice of *σ* and *τ* is also provided in Appendix A.9 and Appendix A.10.

The same kernel also specifies the spectral ordering used by SCI. This ordering is clearest in the idealized translation-invariant setting on *L*^2^(ℝ^*CL*^, d**x**), where the kernel integral operator is convolution by the Gaussian in Eq. (3). Let

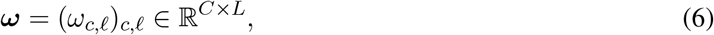

denote a frequency tensor on the flattened history space. The entry *ω*_*c,l*_ is the Fourier frequency associated with channel *c* at lag *l*. The corresponding generalized eigenfunctions are plane waves

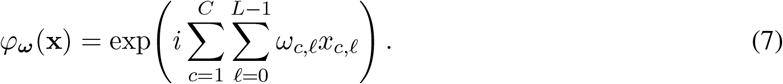

Real-valued spectral features may equivalently be written as the sine and cosine components of the same phase:

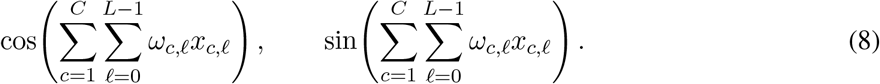

These functions are generalized Fourier eigenfunctions under Lebesgue measure; the empirical eigenvectors used for SCI are defined later under the stimulation distribution *ρ*. The corresponding generalized eigenvalue is (see Appendix A for detailed theoretical calculations.)

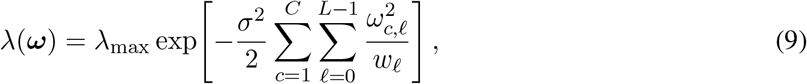

where *λ*_max_ = *λ*(**0**). For the kernel in Eq. (3),

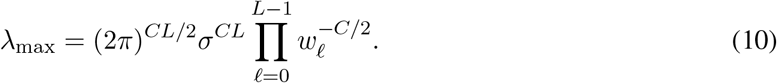

The absolute scale of *λ*_max_ is not used by SCI. The relevant object is the relative spectral decay

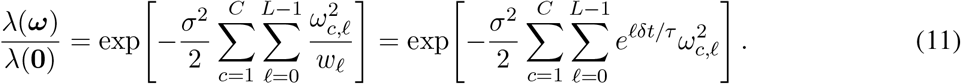

Equation (11) makes the temporal bias explicit. A fixed frequency magnitude at an older lag receives a smaller eigenvalue because 1*/w*_*l*_ = *e*^*lδt/τ*^ grows with *l*. Conversely, for a fixed attenuation level *α* (0, 1), a single-coordinate dependence at lag *l* satisfies

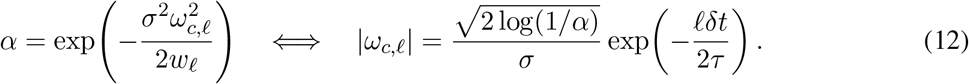

Thus the kernel allows higher-frequency response dependence on recent presynaptic events at the same spectral cost, while older events are represented more coarsely unless the response function uses lower- eigenvalue modes.

In finite simulations, the input histories are not distributed according to Lebesgue measure. They are sampled from the stimulation-induced distribution *ρ*, and the eigenfunctions are data-adapted Mercer modes rather than literal plane waves. The Fourier calculation above is used only to expose the spectral bias of the explicit kernel: early modes correspond to smoother and coarser input dependencies under the chosen metric, whereas later modes correspond to finer spatiotemporal distinctions. The formal *L*^2^(*ρ*) construction and its empirical estimator are given next.

### 3.3 Kernel spectrum and response energy

Let *T*_*ρ*_ be the kernel integral operator on *L*^2^(*ρ*),

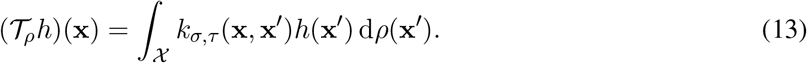

Because the Gaussian kernel is bounded, symmetric, and positive definite, *T* _*ρ*_ is a compact, self-adjoint, positive operator. Its nonzero spectrum admits eigenpairs

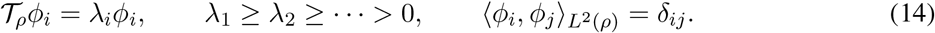

The response function is centered as

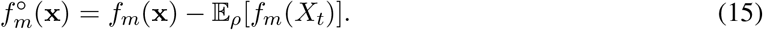

Let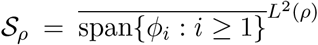 be the nonzero spectral subspace of *Tρ*, and let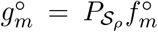 be the *L*^2^(*ρ*)-orthogonal projection of the centered response onto this subspace. The ordered spectral coefficients

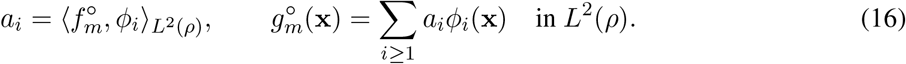

If 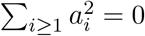, we set SCI_*θ*_(*m*) = 0. Otherwise the cumulative response energy explained by the first *K* kernel modes is

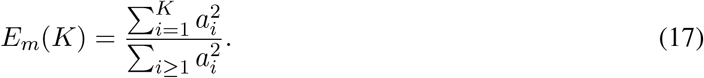

For a target fraction *θ* ∈ (0, 1), the Spectral Complexity Index is

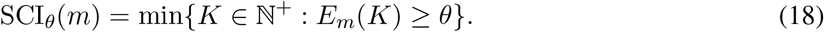

This definition has a direct low-dimensional spectral approximation interpretation. Let

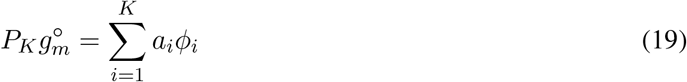

be the orthogonal projection of 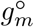 onto the first *K* kernel modes. Then

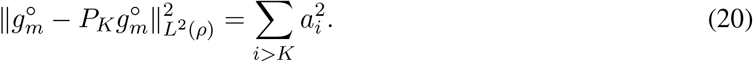

Thus SCI_*θ*_ is the minimum dimension of a low-index spectral subspace required to explain *θ* of the represented centered response energy. The main analysis uses *θ* = 0.8. A low SCI indicates that most response energy is captured by early kernel modes, corresponding to broad aggregate input features under the specified kernel. A high SCI indicates that the response function requires later modes and therefore finer spatiotemporal distinctions in presynaptic history.

### 3.4 Empirical estimator

Given *M* sampled histories and responses 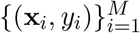, define the Gram matrix

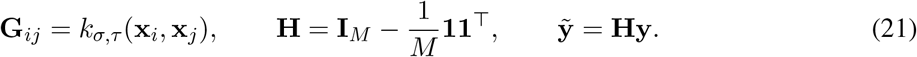

The main definition centers the response but leaves the kernel matrix uncentered: 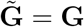. This preserves the analytically tractable spectrum of the uncentered Gaussian kernel. A centered-kernel variant, 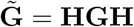evaluated as a robustness control in Appendix C.7.

Let

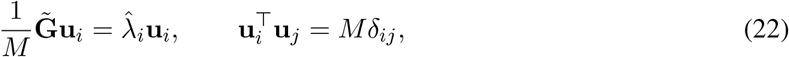

with eigenvectors sorted by decreasing 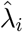. The empirical spectral coefficients are

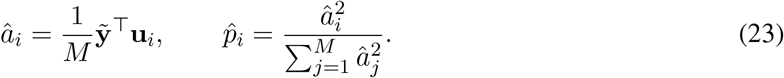

The empirical cumulative energy and SCI are

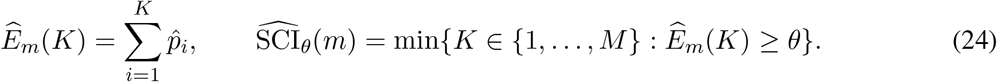

All statistics in the paper are computed at the cell level. Multiple random seeds are used to estimate sampling variability for each cell, and cell-level medians are used for species-level comparisons.

## 4 Experimental setup

### 4.1 Neuron models and stimulation

We evaluate SCI on 35 biophysically detailed cortical neuron models obtained from the Allen Cell Types Database (Allen Institute for Brain Science) using the Allen Software Development Kit (Allen SDK) and simulated in NEURON [17–20]. The analyzed set contains 16 human and 19 mouse cortical neuron models, which consist of 12 spiny neurons for each species, one human sparsely-spiny neuron, and the remaining models (3 human and 7 mouse) are aspiny neurons. Excitatory AMPA+NMDA synapses and inhibitory GABA_*A*_ synapses are distributed over dendritic compartments following the stimulation protocol used for DNN-based functional-complexity analyses [10, 11]. Each simulation contains an initial 500 ms no-input period followed by 10 s of stochastic synaptic drive. Input rates are calibrated so that the output firing rate lies in an approximately 1 Hz regime, reducing trivial excitability confounds.

The SCI input representation uses raw presynaptic event histories, not conductance-weighted currents. This choice makes the input geometry independent of the biophysical response mechanism: differences in synaptic conductance, dendritic morphology, and membrane nonlinearity affect the output function rather than the definition of the input vector. The main response readout is somatic voltage, which includes both subthreshold fluctuations and suprathreshold spike events; firing rates are matched across cells to reduce trivial differences in spike frequency.

To clarify the data generation process and computational efficiency, the *M* evaluated histories do not represent independent 10-second simulations. Instead, we perform 1000 independent 10.5-second simulations per cell. After discarding the initial 500 ms no-input period, sliding windows of length *L* = 200 ms are extracted from the remaining 10-second traces using a Δ_stride_ = 50 ms stride to prevent excessive sample overlap. For each random seed, we uniformly draw *M* = 16384 such history windows from this pool to construct the Gram matrix. Exact eigendecomposition of this *M* ×*M* matrix takes only seconds on a modern GPU. Unless stated otherwise, the main experiments use *δt* = 1 ms, normalized electrotonic distance, *σ* = 1, *τ* = 20 ms, and *θ* = 0.8.

### 4.2 Comparisons and statistics

Species-level comparisons are performed separately for aspiny/sparsely-spiny and spiny groups. The independent statistical unit is the neuron model; time points and random seeds are not treated as independent biological replicates. We report Kolmogorov–Smirnov tests and Welch’s *t*-tests for species comparisons, Spearman rank correlations for agreement with FCI and robustness analyses, and pairwise rank-regression *R* values for morphology analyses. The main external validation compares SCI ranks with FCI ranks on all 35 cells. The method for calculating FCI can be found in Appendix B.1.2. Morphological analyses use standard dendritic summaries and branching descriptors, including total dendritic area, total dendritic length, bifurcation-related branch lengths, terminal branch lengths, and topological features [21, 22].

## 5 Results

### 5.1 SCI separates human and mouse cortical neuron models

**Figure 2.**
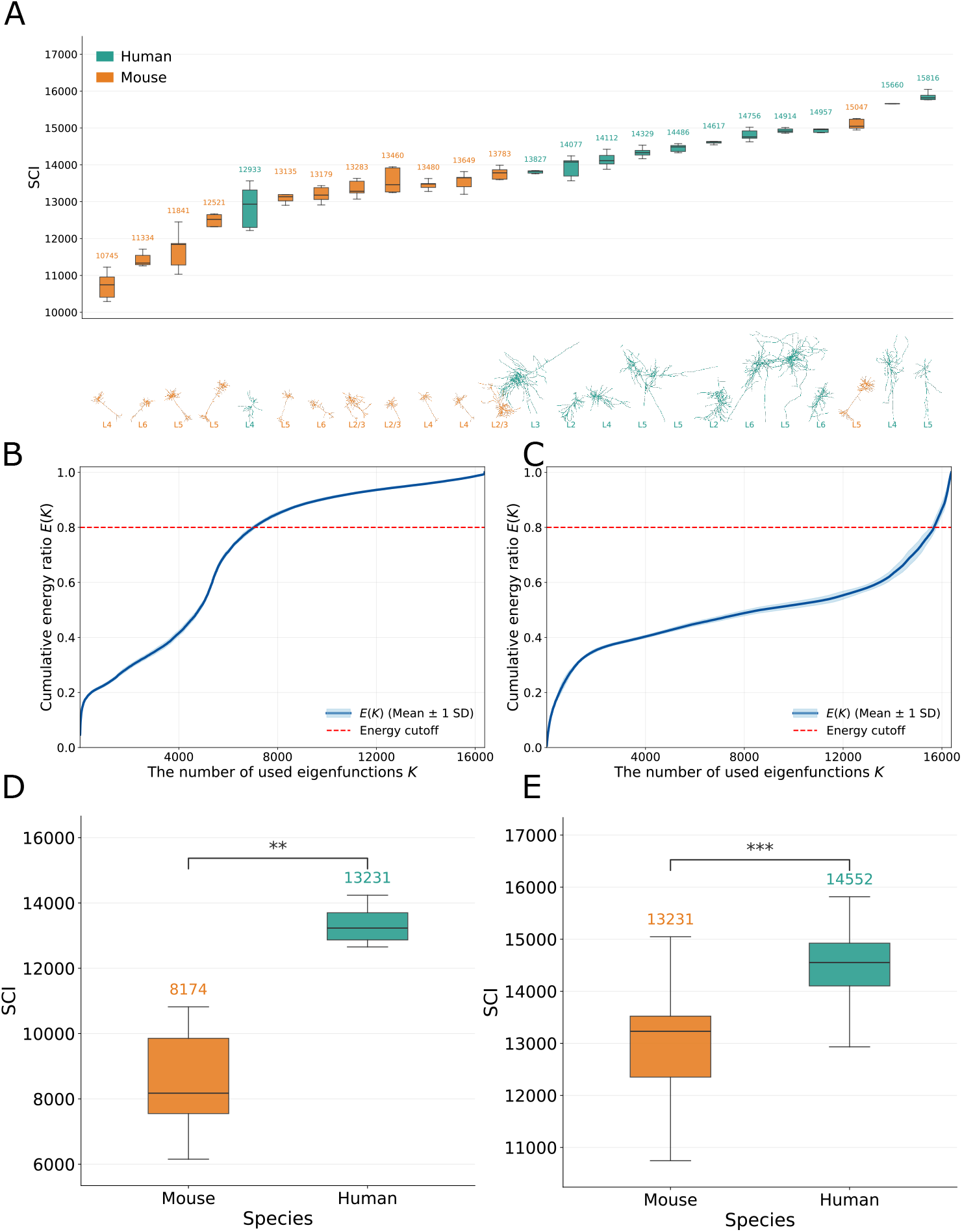
SCI separates human and mouse cortical neuron models. **A**, Spiny cells ranked by cell-level median 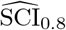. Boxplots summarize random-seed variability for each model (each box summarizes 5 SCI estimates from different window-sampling seeds), with morphology thumbnails and cortical-layer labels shown below. Human models are shown in teal and mouse models in orange. **B–C**, Representative cumulative response-energy curves for lower- and higher-SCI cells. The Model IDs are 480056382 and 626170484 respectively; see Appendix B.2. The thick solid line represents the mean of *E*(*K*), and the shaded area indicates± 1 standard deviation. **D–E**, Species-level summaries using model medians for aspiny/sparsely-spiny cells and spiny cells, respectively. Statistical tests are performed across models rather than across random seeds, with each neuron using its median as a single sample point. Asterisks denote statistical significance according to the Kolmogorov–Smirnov (K–S) test (** *p <* 0.01, *** *p <* 0.001).

SCI identifies species-level differences in both aspiny and spiny groups. Representative cumulative energy curves show the expected spectral signature: a low-SCI mouse neuron reaches the *θ* = 0.8 spectral-energy cutoff using earlier kernel modes, whereas a high-SCI human neuron requires substantially deeper expansion into the spectrum. At the population level, aspiny/sparsely-spiny human neurons have higher SCI than aspiny mouse neurons (K–S *D* = 1.00, *p* = 0.006; Welch *t* = −6.56, *p* = 1.29 ×10^*−*4^; Cohen’s *d* = 3.32; Cliff’s delta = 1.0). The spiny group shows the same direction (K–S *D* = 0.833, *p* = 2.0 ×10^*−*4^; Welch *t* = −3.88, *p* = 9.8× 10^*−*4^; Cohen’s *d* = 1.59; Cliff’s delta = 0.764). The spiny distribution also contains high-SCI mouse cells, indicating that SCI is a cell-level response-function measure rather than a species-label proxy. The results for the aspiny/sparsely-spiny group can be found in Appendix C.1.

### 5.2 SCI agrees with FCI without surrogate-network training

SCI strongly agrees with the DNN-based Functional Complexity Index. Across *n* = 35 neurons, the Spearman rank correlation between SCI rank and FCI rank is *ρ* = 0.797 with *p* = 1.0 ×10^*−*8^. Subgroup correlations remain positive for aspiny/sparsely-spiny cells (*ρ* = 0.700, *p* = 0.016, *n* = 11), spiny cells (*ρ* = 0.672, *p* = 3.2 ×10^*−*4^, *n* = 24), mouse cells (*ρ* = 0.851, *p* = 3.9 ×10^*−*6^, *n* = 19), and human cells (*ρ* = 0.568, *p* = 0.022, *n* = 16). The subgroup results show that the agreement is not solely a species-label effect: SCI aligns with FCI across species and dendrite-type strata while being computed from a different object, the eigenspectrum of an explicit kernel on presynaptic histories.

**Figure 3.**
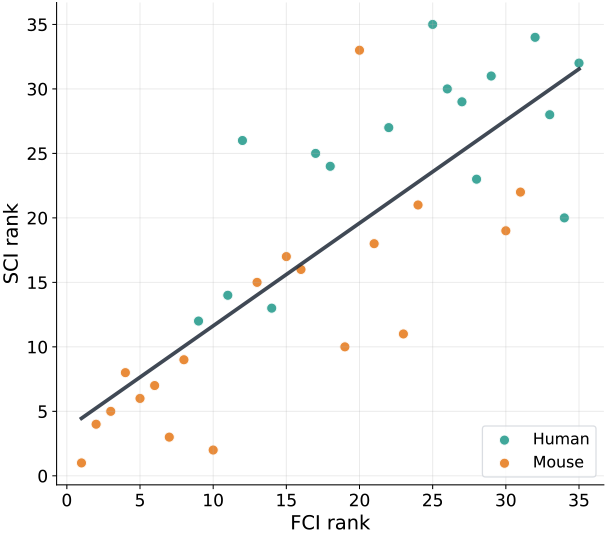
Agreement with FCI. SCI ranks strongly correlate with FCI ranks across all cells (Spearman *ρ* = 0.797, *p* = 1.0 × 10^*−*8^, *n* = 35); colors indicate species.

### 5.3 SCI rankings are stable across kernel and sampling choices

SCI rankings are stable across a broad set of analysis choices. Reducing the sample size to *M* = 4096, 8192, or 12288 preserves high rank agreement with the baseline; for example, *M* = 8192 gives Spearman *ρ* = 0.989. Varying the history window, changing the sampling stride, centering the kernel matrix, or using unnormalized electrotonic distance similarly leaves the ranking nearly unchanged; centered-kernel SCI gives Spearman *ρ* = 0.995. Spectral-energy cutoffs *θ*∈ {0.7, 0.75, 0.85, 0.9}are stable; for example, *θ* = 0.9 gives Spearman *ρ* = 0.973. Kernel parameters show the expected behavior: moderate changes in *σ* and *τ* preserve rankings, whereas an overly narrow bandwidth or excessively long temporal decay scale changes the spectral geometry and correspondingly changes SCI ranks. These boundary cases are consistent with the role of the kernel parameters rather than evidence of arbitrary instability. It should be particularly emphasized that the *L* = 500 ms robustness test supports the working approximation *Y*_*t*_ ≈ *f*_*m*_(*X*_*t*_), as incorporating a longer historical context yields almost unchanged cell-level SCI ranks. See more details in Appendix C.7.

### 5.4 NMDA ablation tests sensitivity to active synaptic nonlinearities

To further isolate the specific contribution of NMDA receptor-mediated nonlinearities to neuronal computational capacity, we calculate SCI across 12 spiny human cortical pyramidal neurons under an NMDA OFF condition. Importantly, input rates are re-calibrated under the NMDA OFF condition to maintain a somatic firing rate around 1 Hz. The deactivation of NMDA receptors yielded a statistically significant reduction in both the mean and median SCI values across the 12 tested spiny human neurons under one-sided Wilcoxon signed-rank tests: for mean SCI, Wilcoxon statistic *V* = 1, one-sided *p* = 5 ×10^*−*4^, Cohen’s *d* = − 1.19; for median SCI, Wilcoxon statistic *V* = 2, one-sided *p* = 7 ×10^*−*4^, Cohen’s *d* = −1.25. Detailed results are reported in Appendix C.4. Notably, despite this significant attenuation in complexity, the SCI of human neurons under the NMDA OFF condition remained persistently higher than the baseline SCI observed in mouse models. Taken together, these results corroborate that while the steep nonlinearity of human NMDA receptors is a critical biological driver of their heightened functional complexity, the other properties of human neurons alone still confer a superior computational advantage over those of mouse neurons.

### 5.5 Morphological correlates of SCI

SCI is related to dendritic morphology but is not fully explained by the measured gross morphological summaries. Pairwise rank regressions using basal morphological features show greater explanatory power in aspiny cells than in spiny cells. In the aspiny cell group, the combination of maximum branch length and total branch number explains a large portion of SCI behavior. In the spiny cell group, however, the same pair of morphological features explains less variance in SCI, suggesting that beyond these coarse morphological summaries, SCI captures intrinsic complexity of neuronal computation. Full morphology heatmaps are reported in Appendix C.5.

### 5.6 Computational cost

The computational cost of SCI is significantly lower than that of existing methods. This reduction comes from two aspects. First and most importantly, robustness tests have demonstrated that computing SCI requires only 100 10-s simulated trajectories, which is far fewer than the 12000 10-second simulated trajectories required by FCI as reported in the original study [11]. Second, after the simulations are completed, the primary overhead of SCI arises solely from computing the *M*× *M* Gram matrix and subsequently performing its eigendecomposition. On a GPU such as the NVIDIA GeForce RTX 5090, the typical computation time for a single seed is around 25 seconds. In contrast, training the TCN (Temporal Convolutional Network) required to compute FCI takes tens of minutes.

## 6 Discussion and limitations

SCI provides a kernel-explicit way to compare single-neuron I/O complexity without fitting a neural-network surrogate. Its main advantage is the explicitness of its assumptions: the stimulation ensemble defines *ρ*, the kernel defines the input geometry, and the eigenspectrum defines the ordering of response components. This makes the resulting complexity axis reproducible and analytically inspectable. In the analyzed cortical neuron models, SCI recovers species-level differences, agrees with FCI, and remains stable across a broad range of implementation choices. Additionally, compared to existing methods, the computation of SCI is significantly faster and consumes fewer resources, providing a more efficient approach for investigating neuronal complexity.

The index is still not fully intrinsic. SCI depends on the input ensemble, the response readout, and the kernel. This dependence is appropriate for comparing neurons under a standardized stimulation protocol, but it limits claims about universal complexity independent of input statistics. The main analysis uses somatic voltage as the response because it provides a dense readout of the finite-history I/O transformation and includes both subthreshold fluctuations and suprathreshold spike events; spike-window outputs are a natural robustness analysis. The finite-history treats the observed voltage as an approximately deterministic function of the chosen history window. The channelization also coarse-grains branch identity by synapse type, dendritic domain, and electrotonic distance. These limitations identify controlled extensions of the framework rather than changes to its core definition.

The present results complement recent work connecting human neuronal specialization to morphology, synaptic physiology, transcriptomics, and connectomics [23–25]. They also complement current NeuroAI efforts that incorporate dendritic mechanisms into artificial architectures [26]. SCI provides a reproducible spectral coordinate system for asking which parts of a neuron’s I/O map depend on broad input summaries and which require finer spatiotemporal distinctions. This coordinate system can be applied to targeted perturbations of synapse type, NMDA conductance, dendritic channel distributions, or branch-aware input representations.

## Acknowledgement

This work was supported by the National Key R&D Program of China (Project No. 2025YFA1016700) and the National Natural Science Foundation of China (Grant No. 62576011).

## A Theoretical details

This appendix gives the spectral calculation underlying SCI. The derivation starts from the one-dimensional Gaussian kernel on Lebesgue measure, then extends to the *d*-dimensional Gaussian kernel, and finally specializes to the time-weighted presynaptic-history kernel used in the main text. The Lebesgue-measure calculation is used to expose the spectral bias of the kernel. The empirical SCI estimator in the main text estimates the corresponding kernel spectrum under the stimulation-induced input distribution.

Throughout this appendix, Gaussian kernels are written without density normalization. Multiplying the kernel by a constant rescales all eigenvalues by the same factor and does not change eigenfunctions, relative spectral decay, or SCI.

### A.1 One-dimensional Gaussian kernel on Lebesgue measure

Let

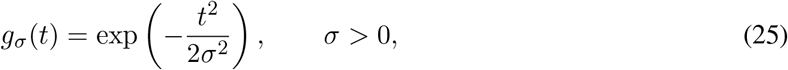

and consider the one-dimensional translation-invariant Gaussian kernel

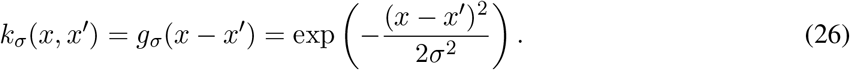

On *L*^2^(ℝ, d*x*), the associated integral operator is

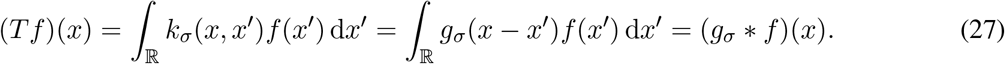

Since *g*_*σ*_ ∈ *L*^1^(ℝ), Young’s inequality gives

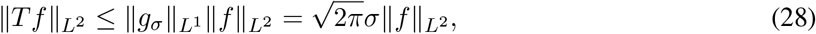

so *T* is a bounded linear operator on *L*^2^(ℝ).

We use the Fourier-transform convention

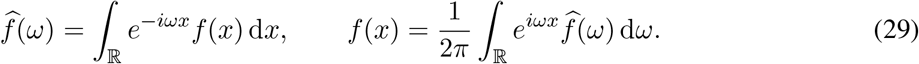

Under this convention,

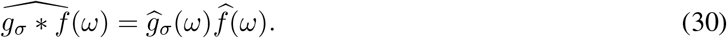

It remains to compute 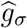. Directly,

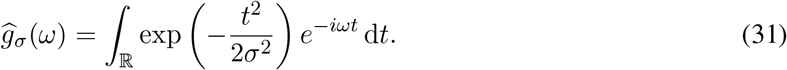

Completing the square,

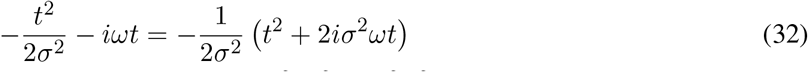

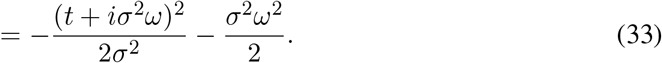

Hence

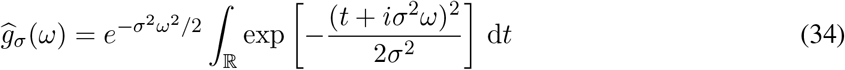

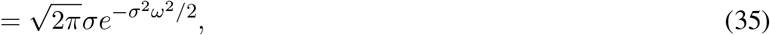

where the final equality follows from the analytic continuation of the Gaussian integral. Define

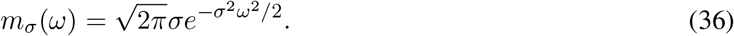

Then

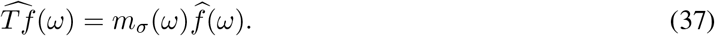

Thus the Fourier transform diagonalizes the Gaussian convolution operator.

The operator in Eq. (27) has no nonzero ordinary *L*^2^(ℝ) eigenfunction. Suppose *Tf* = *λf* for some *f* ∈ *L*^2^(ℝ). Taking Fourier transforms gives

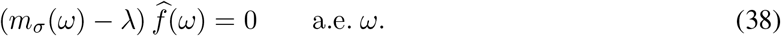

Therefore 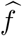 can be supported only on the level set

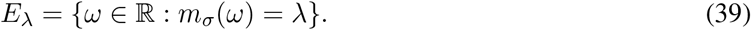

The multiplier satisfies

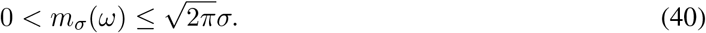

If 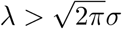, then *E*_*λ*_ = ∅. If *λ* =2*πσ*, then 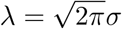. If 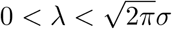, then

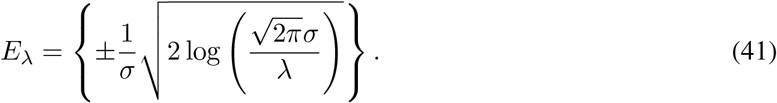

In every case, *E*_*λ*_ has Lebesgue measure zero. Since 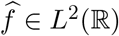, support on a null set implies 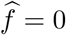 almost everywhere, hence *f* = 0. For *λ* = 0, the multiplier is strictly positive for every finite *ω*, so again there is no nonzero eigenfunction. Consequently,

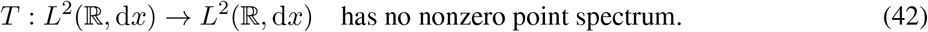

As a multiplication operator in the Fourier domain, its spectrum is the closure of the essential range of *m*_*σ*_:

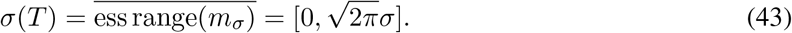

This is continuous spectrum.

Although ordinary *L*^2^ eigenfunctions do not exist, plane waves are generalized eigenfunctions. For *ϕ*_*ω*_(*x*) = *e*^*iωx*^,

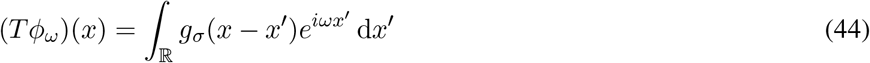

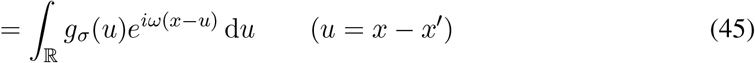

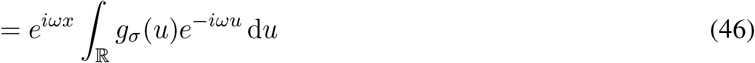

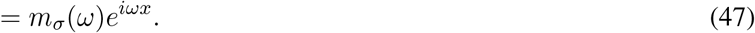

Hence, in the generalized Fourier sense,

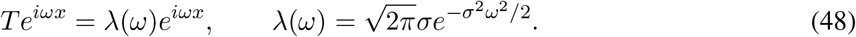

With the *δ*-normalized plane wave

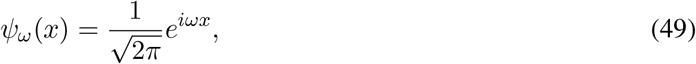

one has

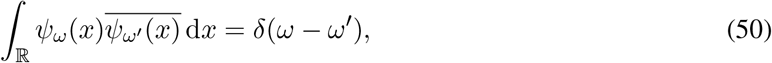

and the kernel admits the continuous Fourier expansion

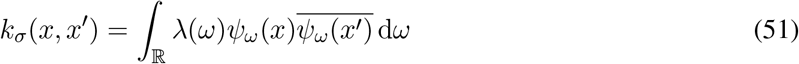

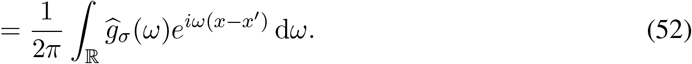

The largest generalized eigenvalue is

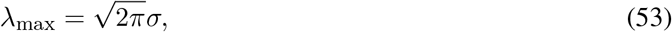

attained at *ω* = 0, corresponding to the constant generalized eigenfunction. For any 0 *< λ < λ*_max_,

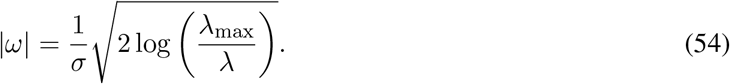

Thus smaller eigenvalues correspond to higher-frequency plane waves. In this idealized one-dimensional calculation, the Gaussian kernel orders functions from low-frequency, smooth dependencies to high-frequency, rapidly varying dependencies.

### A.2 *d*-dimensional isotropic Gaussian kernel

Let **x, x**^*′*^ ∈ ℝ^*d*^ and

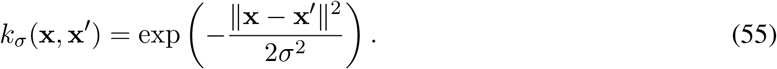

Writing **z** = **x** − **x**^*′*^, define

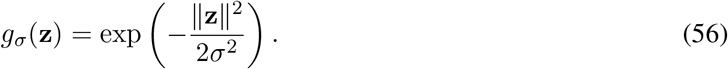

The integral operator on *L*^2^(ℝ^*d*^, d*x*) is again convolution:

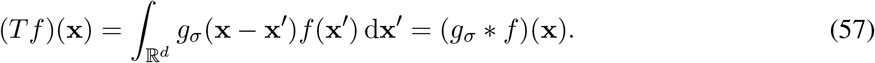

We use

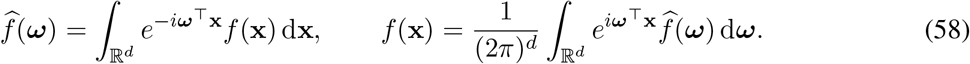

Since

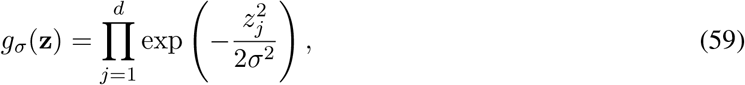

Fubini’s theorem and the one-dimensional calculation yield

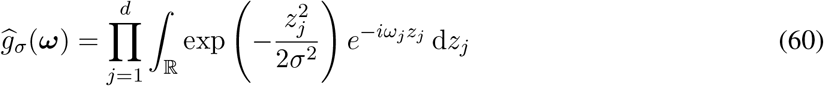

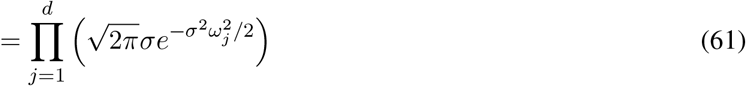

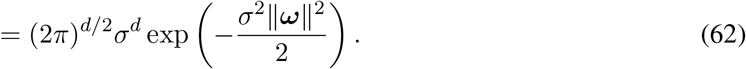

Define

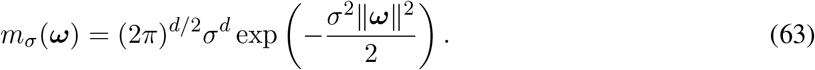

Then

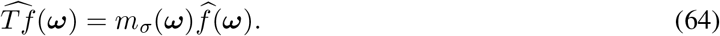

The same level-set argument shows that *T* has no nonzero ordinary *L*^2^(ℝ^*d*^) eigenfunction. If *Tf* = *λf*, then 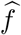 is supported on

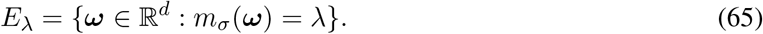

The maximum value is

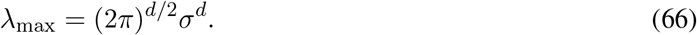

For 0 *< λ < λ*_max_, *E*_*λ*_ is the sphere

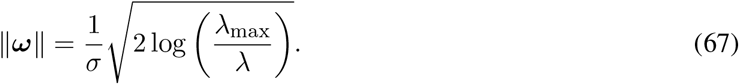

This sphere has *d*-dimensional Lebesgue measure zero. Hence no nonzero *L*^2^ eigenfunction exists. The spectrum is

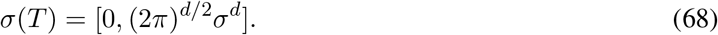

For each ***ω*** ∈ ℝ, the plane wave

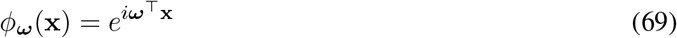

is a generalized eigenfunction:

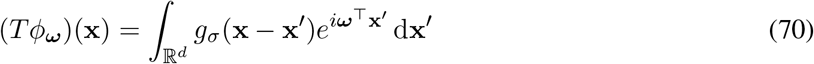

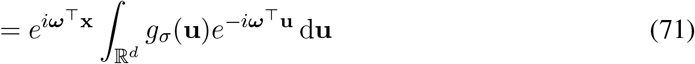

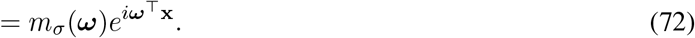

Thus

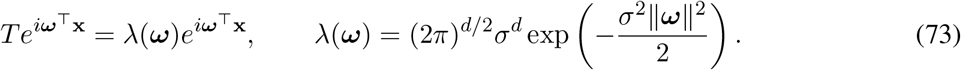

With

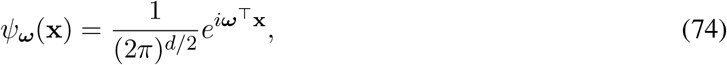

the kernel expansion is

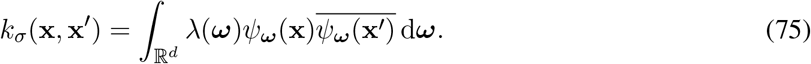

Equation (67) shows the degeneracy of the isotropic Gaussian spectrum. In dimensions *d* ≥ 2, a fixed eigenvalue corresponds to an entire frequency sphere. Real-valued generalized eigenfunctions can be taken as

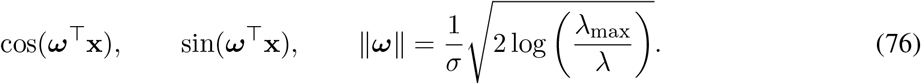

The eigenvalue depends on the frequency magnitude, not on the frequency direction.

### A.3 The time-weighted presynaptic-history kernel

The kernel used by SCI is a Gaussian kernel on channelized presynaptic histories. A history is indexed by channel *c* = 1, …, *C* and lag *l* = 0, …, *L* − 1, and is flattened as a vector in ℝ^*CL*^. The kernel is

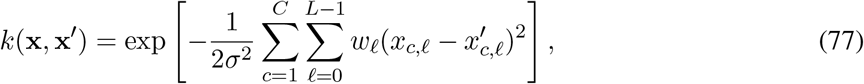

where

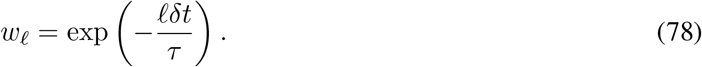

The exponent is separable across all coordinates, so

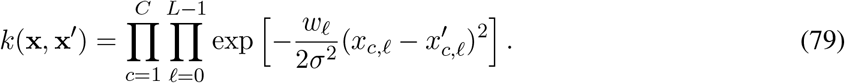

For a fixed lag *l*, the coordinate-wise kernel has length scale

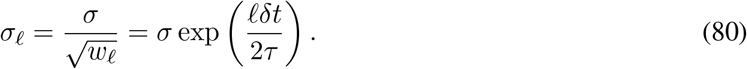

Thus older input lags are compared with a larger length scale.

Let *p* = *CL* and *z* = *x* − *x*^*′*^. Define

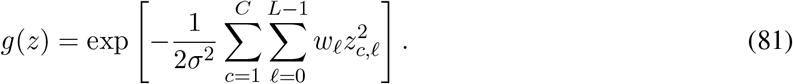

On *L*^2^(ℝ^*CL*^, d*x*), the integral operator is

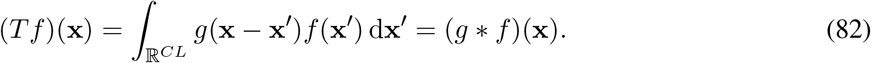

The Fourier transform of *g* is computed coordinate by coordinate:

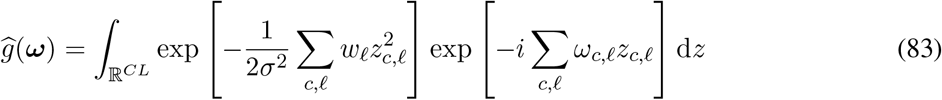

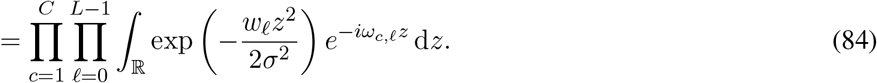

For a fixed *l*, let *a*_*l*_ = *w*_*l*_*/σ*^2^. The one-dimensional Gaussian integral gives

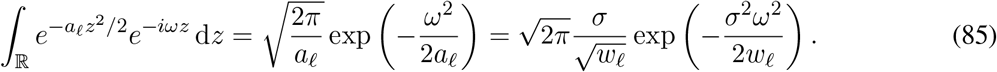

Combining all factors,

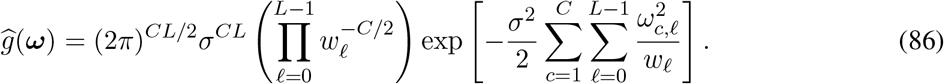

Substituting *w*_*l*_ = *e*^*−lδt/τ*^ yields

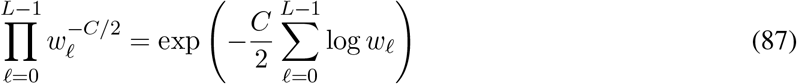

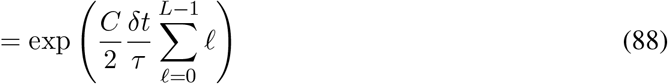

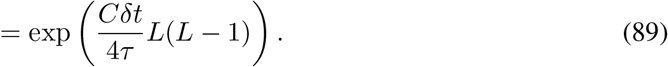

The maximum generalized eigenvalue is therefore

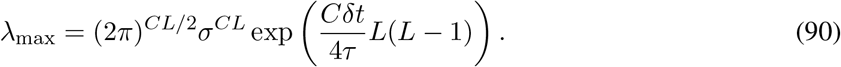

The relative eigenvalue is the dimensionless quantity

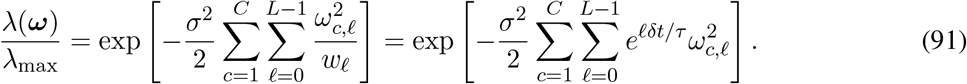

This is the spectral ordering used to interpret SCI.

The generalized eigenfunctions remain plane waves on the flattened history space:

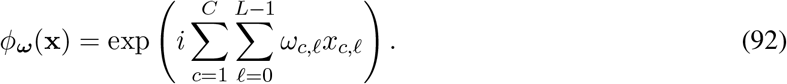

They satisfy

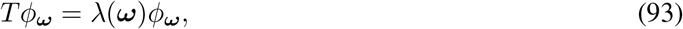

Where

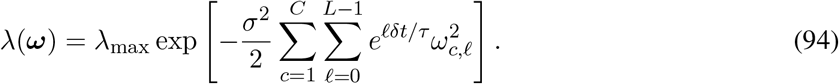

As before, these are generalized eigenfunctions under Lebesgue measure, not ordinary *L*^2^(ℝ^*CL*^) eigenfunctions. The spectrum is continuous:

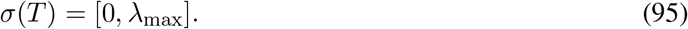

For a fixed eigenvalue 0 *< λ* ≤ *λ*_max_, define

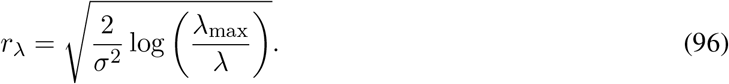

Then

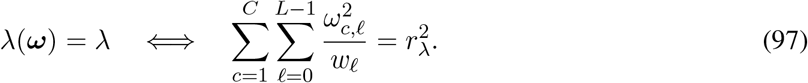

Thus a fixed eigenvalue corresponds to a frequency ellipsoid. The real-valued generalized eigenfunctions associated with this ellipsoid can be written as

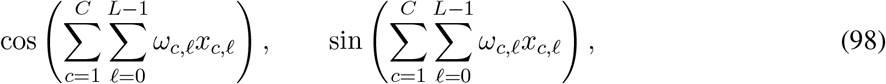

with ***ω*** satisfying Eq. (97).

Equation (91) makes the temporal inductive bias explicit. The penalty on *ω*_*c,l*_ is multiplied by *e*^*lδt/τ*^. For the same frequency magnitude, dependence on older lags is assigned a smaller eigenvalue than dependence on recent lags. Equivalently, if only one coordinate at lag *l* is active, then the frequency that gives a fixed attenuation ratio *α* ∈ (0, 1) satisfies

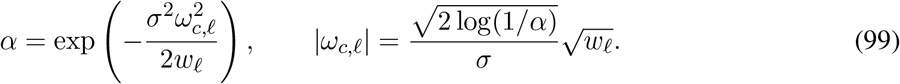

The effective bandwidth at lag *l* therefore scales as

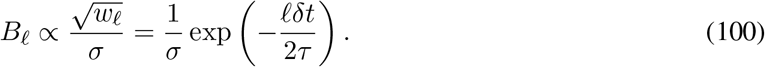

Recent inputs support higher-frequency dependencies at a given spectral cost; older inputs are represented more coarsely unless the response function uses low-eigenvalue modes.

## A.4 From Lebesgue spectrum to the stimulation distribution

The Fourier calculations above describe the spectral bias of the uncentered Gaussian kernel under Lebesgue measure. SCI itself is defined under the input distribution *ρ* induced by the stimulation protocol. Let

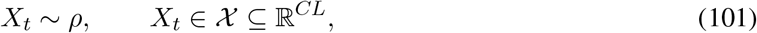

and let *Y*_*t*_ be the scalar response used for analysis. In the main experiments, *Y*_*t*_ is the next-bin somatic membrane voltage. The input–output map is represented by the finite-history regression function

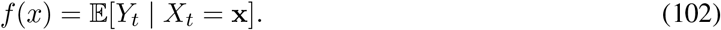

The empirical estimator uses observed voltage samples as proxies for this finite-history map, corresponding to the working approximation

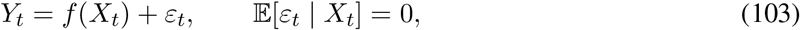

with residual energy treated as small for the calibrated voltage readout and chosen history window. This approximation is supported operationally by the history-window robustness in Appendix C.7, where *L* = 500 ms gives Spearman *ρ* = 0.997 relative to the *L* = 200 ms baseline.

Define the kernel integral operator on *L*^2^(*ρ*) by

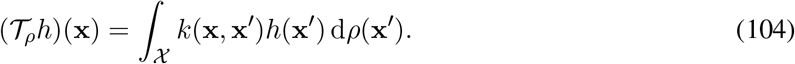

The Gaussian kernel is bounded, symmetric, and positive definite. Since *ρ* is a probability measure,

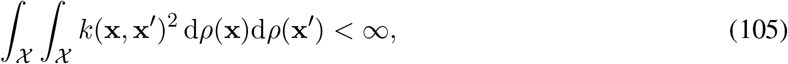

so *T*_*ρ*_ is Hilbert–Schmidt and hence compact. Symmetry of *k* gives self-adjointness, and positive definiteness gives (we work with real-valued *L*^2^(*ρ*) functions)

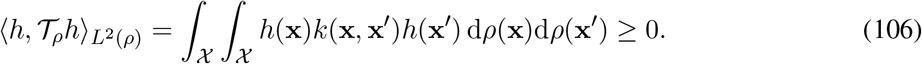

By the spectral theorem for compact self-adjoint operators, the nonzero spectrum is discrete and admits eigenpairs

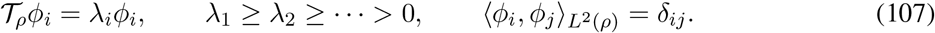

Under the usual Mercer conditions, this yields the standard kernel expansion; more generally the expansion is understood in *L*^2^(*ρ* ×*ρ*) sense [16, 27].

The centered response function is

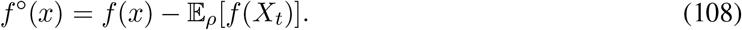

SCI projects Then *f*^*°*^ onto the nonzero spectral subspace of T_*ρ*_. Let 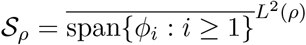 and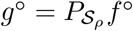 Then

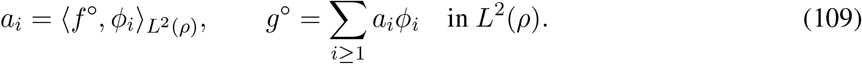

If 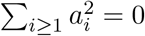, we set SCI_*θ*_(*f, ρ, k*) = 0. Otherwise, the cumulative response energy explained by the first *K* modes is

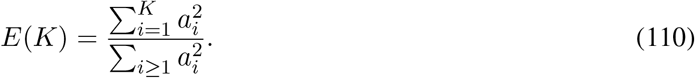

For *θ* ∈ (0, 1), the Spectral Complexity Index is

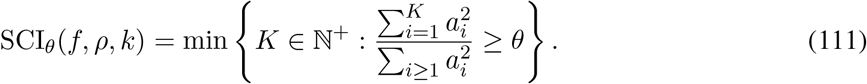

The notation SCI_*θ*_(*f, ρ, k*) emphasizes that SCI is an operational complexity of a response function under a specified stimulation ensemble and an explicit input geometry. It does not depend on a trained neural-network surrogate. It does depend on the kernel and input distribution.

If an eigenvalue has multiplicity greater than one, the total energy assigned to the whole eigenspace is invariant, while individual coefficients and the exact within-cluster mode count can depend on the chosen orthonormal basis. SCI uses a fixed within-eigenspace ordering as a mode-counting convention. The empirical SCI reported in this paper is defined directly by the finite-sample eigendecomposition and does not aggregate degenerate or nearly degenerate clusters.

### A.5 Response centering and kernel centering

The main estimator centers the response and uses the uncentered kernel. Response centering removes the input-independent DC component:

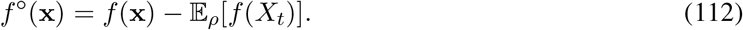

A constant response has zero centered energy and contributes no SCI.

Centering the kernel would replace *k* by the *ρ*-dependent centered kernel

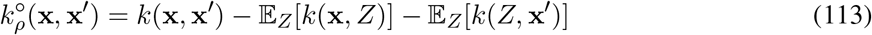

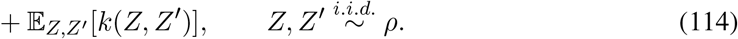

For a generic stimulation distribution *ρ*, 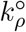 is not translation invariant even when *k* is a Gaussian kernel. The uncentered Gaussian kernel is therefore used as the primary definition, because its spectral bias is the Fourier structure derived in Appendix A.1–A.3. The centered-kernel version is a robustness control in which the same response-energy definition is applied to the empirical centered Gram matrix.

### A.6 Finite-sample estimator

Given samples 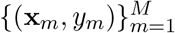, define

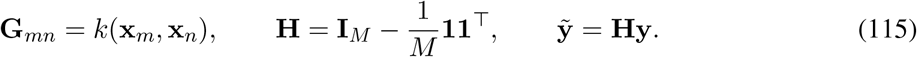

The empirical measure is

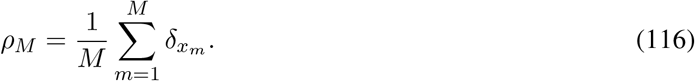

The finite-sample operator associated with *ρ*_*M*_ has sample values satisfying the Gram eigenproblem

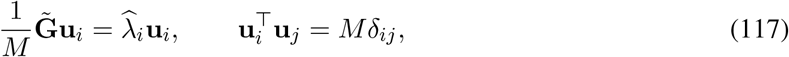

where 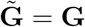for the main estimator and 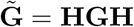 for the centered-kernel robustness estimator.

The normalization 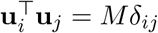 makes the vectors orthonormal under the empirical inner product

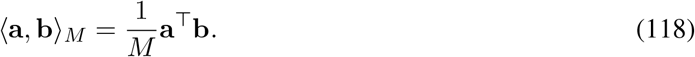

The empirical spectral coefficient of the centered response is

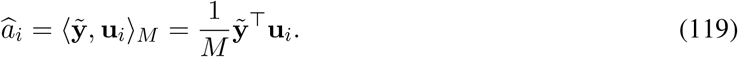

If 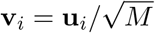 is Euclidean-normalized, then the same coefficient is

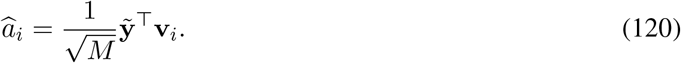

By Parseval’s identity in the empirical inner product, when the full eigenbasis is used,

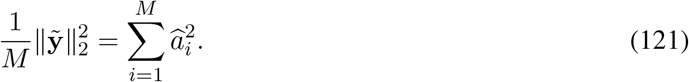

The empirical energy distribution is

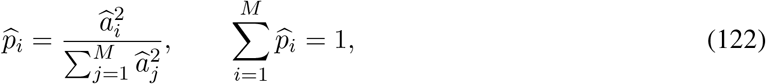

and

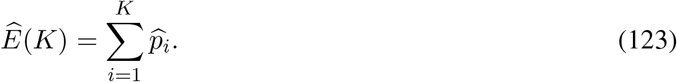

The empirical SCI is

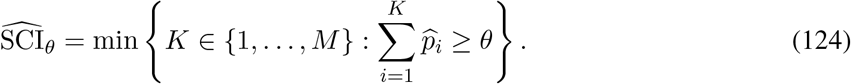

Numerically, all *M* eigenvectors returned by the eigendecomposition are kept and sorted by decreasing eigenvalue; no zero- or near-zero-eigenvalue thresholding and no degenerate-cluster aggregation is applied. The response vector is centered in every analysis.

### A.7 Relation to RKHS smoothness

The spectral ordering also connects SCI to the smoothness bias of the Gaussian RKHS. For a translationinvariant kernel *k*(**x, x**^*′*^) = *g*(**x** − **x**^*′*^) on ℝ^*p*^, let *m*(***ω***) = *ĝ* (***ω***). Under the Fourier convention used above, the RKHS norm of a function *f* with Fourier transform 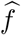 is

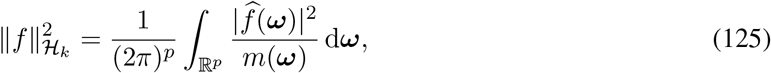

when the integral is finite. For the time-weighted kernel,

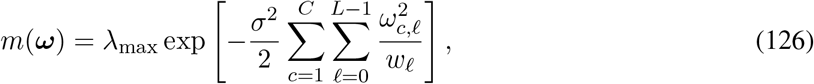

so

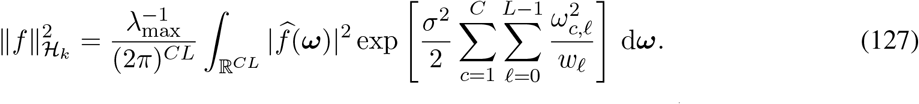

High weighted frequencies incur exponentially larger RKHS cost. Since 1*/w*_*l*_ = *e*^*lδt/τ*^, fine dependence on older input lags is especially costly under this kernel. SCI does not use the RKHS norm directly. It uses the same spectral ordering but measures the number of modes needed to explain a fixed fraction of unweighted *L*^2^(*ρ*) response energy. This avoids excessive sensitivity to the smallest empirical eigenvalues while retaining the smooth-to-fine ordering specified by the kernel.

### A.8 Interpretation of spectral modes

In the Lebesgue–Fourier calculation, a generalized mode has the form

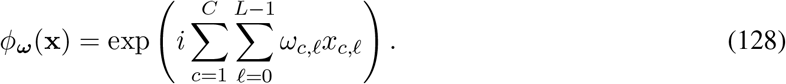

The vector ***ω*** specifies a spatiotemporal sensitivity pattern over channels and lags. A large coefficient *ω*_*c,l*_ means that the mode changes rapidly with the input coordinate *x*_*c,l*_; a zero coefficient means that the mode is insensitive to that coordinate. Multiple nonzero entries correspond to joint dependence across several channels and time lags.

The maximum-eigenvalue mode corresponds to ***ω*** = **0**, hence to the constant function. After response centering, this DC response component is removed. Large eigenvalues correspond to small weighted frequency

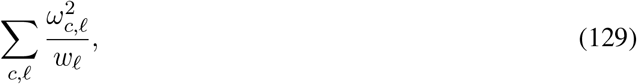

and therefore to broad aggregate input features. These modes can represent total input drive, smooth E/I balance, broad basal or apical input trends, and low-order smooth response components. For small *ω*,

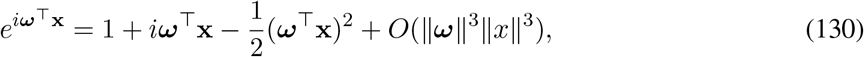

so low-frequency modes naturally include constant, linear, and low-order smooth interactions.

Intermediate eigenvalues correspond to moderate spatiotemporal resolution. They can represent contrasts between channels, coarse synchrony within a time window, or low-order nonlinear interactions across nearby dendritic domains. For example, if 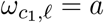 and 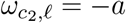, then the mode depends on 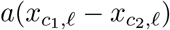, an input contrast between two channels. If *ω*_*c,l*_ has the same sign across many channels, the mode depends on an aggregate population drive.

Small eigenvalues correspond to large weighted frequencies and hence to response components that vary sharply over the input-history space. These modes are needed for fine spike-timing sensitivity, local coincidence detection, input-order selectivity, branch-specific clustered input effects, threshold-like transitions, and other combinatorial nonlinearities. A high SCI indicates that a substantial fraction of the centered response energy is not captured by broad aggregate modes and must be represented by later, finer kernel modes.

In empirical data, eigenfunctions are not literal Fourier plane waves. They are data-adapted eigenfunctions of *T* _*ρ*_. The Fourier calculation specifies the kernel’s ordering principle, while the stimulation distribution adapts the actual eigenfunctions to the sampled input regime. SCI uses this ordered empirical spectrum to measure how far into the kernel-defined hierarchy the neuronal response function extends.

### A.9 Choice of *σ*

Consider

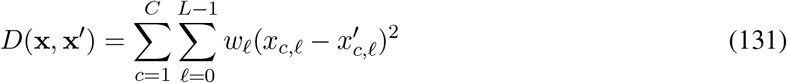

In our framework, if *x*_*c,l*_ and 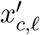 differ by a single spike in the most recent time bin (*l* = 0), then *w*_*l*_ = 1 and *D* changes by 1. With *σ* = 1, this produces a kernel attenuation factor exp(−1*/*2) ≈0.607, so a single recent-bin spike difference remains resolvable. Empirically, the medians of the upper triangular part of *D* excluding the diagonal mostly lie around [20, 60]. Taking *D* = 40 as a representative value, *σ* = 1 gives a typical off-diagonal Gram entry exp(−40*/*2) ≈ 2.1 ×10^*−*9^, whereas *σ* = 0.5 gives exp[−40*/*(2 ×0.5^2^)] ≈ 1.8 ×10^*−*35^. The issue for *σ* = 0.5 is that almost all off-diagonal entries become negligible relative to the diagonal entries, limited by the machine epsilon of float64. The Gram matrix then becomes close to the identity, the empirical spectrum becomes nearly flat, and the eigenvector ordering is dominated by small sampling and numerical perturbations. Increasing *σ* broadens the kernel and reduces single-spike resolution. Thus *σ* = 1 is a heuristic compromise between recent-bin event resolution and a nondegenerate empirical spectral geometry; Appendix C.7 shows that SCI rankings remain robust to broader choices *σ* = 1.5 and *σ* = 2.0, while the overly narrow *σ* = 0.5 boundary case changes the ordering.

### A.10 Choice of *τ*

The parameter *τ* sets the recency bias of the temporal metric. In cortical pyramidal neurons, the physiological membrane time constant is typically around 10–30 ms [7, 8]. Setting *τ* = 20 ms is consistent with this physiological range. Mathematically, a larger *τ* slows the decay of *w*_*l*_ = exp(−*lδt/τ*) and therefore assigns more metric weight to older lags; equivalently, the effective length scale 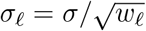 grows more slowly with lag. Thus *τ* = 50 ms does not smooth away older inputs. Rather, it weakens the intended recency bias and makes early spectral modes spend resolution on long-past input variation that may lie outside the dominant physiological integration window. This changes the kernel geometry, consistent with the lower rank agreement observed for the *τ* = 50 ms boundary case in Appendix C.7. Moderate perturbations, such as *τ* = 10 ms and *τ* = 30 ms, preserve the ranking.

## B Experimental details

### B.1 Simulation details

#### B.1.1 SCI calculation details

All simulations use the same pipeline before any SCI-specific window sampling. For each cell, the Allen biophysical model is loaded in NEURON, and synaptic input is placed only on basal and apical dendritic compartments. Each dendritic or apical segment receives one excitatory AMPA–NMDA point process and one inhibitory GABA_*A*_ point process. A segment *r* of length *l*_*r*_ *µ*m is assigned ⌈ *l*_*r*_⌉ source-level presynaptic event channels for each sign; these source-level channels are mapped to the segment-level point process by a model-level sparse gather matrix. The nonzero gather weights are sampled once per model from Unif(0, 1) and then held fixed across all trials of that model, so different trials change the presynaptic event times but not the segment-level synaptic layout. The NMDA-ablation condition used the otherwise identical simulation pipeline with the NMDA conductance ratio set to zero.

Each trial contains 500 ms of initial silence followed by 10 s of stochastic synaptic drive. Source-level excitatory and inhibitory event trains are generated on a 0.1 ms time grid. For each trial, a coarse time-varying event-rate trace is sampled by drawing a rate-update interval from a discrete set between 25 ms and 450 ms, adding integer jitter of ±20 ms. For each sign, the piecewise-constant rates are sampled from zero up to a sign-specific ceiling measured in spikes per synapse per 100 ms, so these ceilings set the amplitude of the coarse pre-smoothing drive rather than directly determining the final event count; in the present study, the excitatory ceiling is adjustable whereas the inhibitory ceiling is held fixed. The result is then smoothed with a normalized Gaussian temporal window. The Gaussian smoothing scale is sampled per trial from values between 25 ms and 600 ms, again with integer jitter ±20 ms; the smoothing-window length is seven times the sampled scale. Binary source-level events are then sampled independently, conditional on the smoothed rate trace by the discrete stochastic sampler implemented in the pipeline. We therefore treat the inputs as stochastic event trains rather than as exact Poisson spike trains.

Source-level events are aggregated to segment-level inputs by left-multiplying the corresponding sparse event matrices by the fixed gather matrix. A nonzero event in a segment-level row is delivered to the appropriate NEURON point process with NetCon weight equal to the baseline synaptic conductance multiplied by the aggregate event weight. Event times are generated on the 0.1 ms input grid and passed to the full compartmental model. The somatic membrane voltage is recorded throughout the trial, interpolated onto the same 0.1 ms grid, and then averaged into 1 ms bins for the arrays used by SCI. Segment-level input arrays are downsampled to 1 ms by summing the ten 0.1 ms source bins in each saved bin. Output spike times are detected only for quality control, using a model-specific somatic voltage threshold estimated from an independent current-clamp run; SCI itself used the saved somatic voltage as the response variable. The downstream SCI channels are formed from saved weighted event counts, so synaptic conductance parameters affected the simulated voltage response but not the metric between input histories.

The retained simulation set is defined by firing-rate quality control after the initial silence period. For every trial, the average post-silence firing rate is computed from threshold crossings of the somatic voltage, and a trial is retained only if this rate lay in [0.8, 1.2] Hz. This filtering is applied separately within each model and data split before coarse graining, HDF5 aggregation, and SCI window sampling.

For the SCI input coordinates, each segment is assigned to an apical or basal dendritic domain and to a bin of electrotonic distance from the soma. Electrotonic distance is computed with the NEURON impedance tool at zero frequency. For a segment *r*, the unnormalized distance is 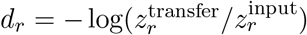, where 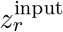 is the local input impedance and 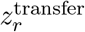 is the transfer impedance from that segment to the soma. The ratio is clipped only to avoid undefined logarithms, and distances are normalized within each cell by dividing by the maximum electrotonic distance over its dendritic and apical segments. In the main analysis, normalized distances are divided into 50 bins per dendritic domain. For each 1 ms time bin, segment-level excitatory and inhibitory events are summed within each combination of sign, dendritic domain, and distance bin, producing 2×2× 50 = 200 input channels ordered as excitatory-apical, excitatory-basal, inhibitory-apical, and inhibitory-basal. A robustness variant used bins of the unnormalized electrotonic distance, with distances beyond the chosen upper range assigned to the final bin.

After trial filtering and coarse graining, all retained training trials for a model are gathered into model- level HDF5 arrays containing the 200 ×*T* coarse-grained input matrix and the aligned somatic voltage trace for each trial. SCI windows are drawn from the post-silence interval only. A candidate window is indexed by its first 1 ms bin *s*≥ 500, contains the coarse-grained input segment from *s* through *s* + *L*− 1, and is paired with the somatic voltage at *s* + *L*. Candidate starts are spaced by Δ_stride_*/*1 ms = 50 bins, and *M* windows are sampled without replacement for each SCI seed, with *M* set to the value reported in Section 4.1 for the main analysis. In the SCI vector, the saved window is reversed so that *l* = 0 corresponds to bin *s* + *L*−1, the most recent input bin. Multiple window-sampling seeds are used for every model, and the reported cell-level value is the median across seeds unless otherwise stated.

For each sampled set, the implementation constructed the pairwise time-weighted squared distance used in Eq. (3). The temporal weights are *w*_*l*_ = exp(− *lδt/τ*), with the most recent bin receiving weight one; in the main analysis *δt* = 1 ms and *τ* = 20 ms. The Gaussian Gram matrix is **G**_*ij*_ = exp[− **D**_*ij*_*/*(2*σ*^2^)] with *σ* = 1. The default SCI calculation used the uncentered Gram matrix and centered only the voltage response, matching Eq. (24). Eigenvectors are ordered by decreasing eigenvalue, voltage coefficients are squared and normalized into a modal energy distribution, and 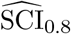 is reported as the first mode index at which the cumulative modal energy reached 0.8.

The centered-kernel robustness control used the same response centering but replaced the Gram matrix by the centered Gram matrix **HGH**. The same spectral ordering is applied to the centered kernel, and 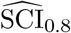 is computed from the resulting eigenvectors and eigenvalues.

#### B.1.2 FCI calculation details

We configured the Temporal Convolutional Network (TCN) and performed the computations using the same method as Aizenbud et al. [11], except that the signal generation adopted the same simulation approach used before calculating the SCI, which differs from the original paper. Additionally, for computational tractability, we used only 1000 10-second simulation trajectories as the training set and 200 10-second simulation trajectories as the test set. To control for data-generation differences, in the main text, we extracted the trajectories required for the SCI calculation from the 1000 simulations within this training set as well. However, the robustness analysis has demonstrated that using 100 trajectories is sufficient to achieve high rank agreement (Spearman *ρ* = 0.992 compared to the baseline setting; see Table 4); therefore, the data volume required for SCI is less than 1*/*10 of that required by currently existing methods.

### B.2 Biophysical model details

All biophysical models of the neurons used in this paper are derived from the Allen SDK [19], all of which are biophysical perisomatic models. Detailed information about the spiny neuron models is provided below (the Index column corresponds to the left-to-right order in Figure 2**A**.):

**Table 2:** Spiny model details.

| Index | Species | Cell ID | Model ID | Brain Region | Layer | Transgenic Line |
|---|---|---|---|---|---|---|
| 1 | mouse | 486893033 | 487245719 | VISp | 4 | Scnn1a-Tg3-Cre |
| 2 | mouse | 471819401 | 478513398 | VISp | 6a | Ntsr1-Cre_GN220 |
| 3 | mouse | 471129934 | 480624414 | VISp | 5 | Rbp4-Cre_KL100 |
| 4 | mouse | 486111903 | 486556811 | VISp | 5 | Rbp4-Cre_KL100 |
| 5 | human | 508405843 | 538906694 | MTG | 4 | NA |
| 6 | mouse | 485909730 | 486509232 | VISp | 5 | Cux2-CreERT2 |
| 7 | mouse | 488680211 | 489932435 | VISp | 6a | Rorb-IRES2-Cre |
| 8 | mouse | 477127614 | 478809991 | VISp | 2/3 | Slc17a6-IRES-Cre |
| 9 | mouse | 485468180 | 485720587 | VISp | 2/3 | Nr5a1-Cre |
| 10 | mouse | 486502127 | 486909496 | VISp | 4 | Rorb-IRES2-Cre |
| 11 | mouse | 488420599 | 488763268 | VISp | 4 | Rbp4-Cre_KL100 |
| 12 | mouse | 487661754 | 488462783 | VISp | 2/3 | Cux2-CreERT2 |
| 13 | human | 548425467 | 626170529 | MFG | 3 | NA |
| 14 | human | 508421103 | 549384713 | MTG | 2 | NA |
| 15 | human | 527952884 | 626170709 | MTG | 4 | NA |
| 16 | human | 528042185 | 560692892 | MTG | 5 | NA |
| 17 | human | 527942865 | 626170718 | MTG | 5 | NA |
| 18 | human | 541563728 | 626170538 | MTG | 2 | NA |
| 19 | human | 548494556 | 626170511 | MFG | 6 | NA |
| 20 | human | 529898751 | 626170619 | ITG | 5 | NA |
| 21 | human | 558211203 | 626170502 | MTG | 6 | NA |
| 22 | mouse | 488679042 | 489932597 | VISp | 5 | Rorb-IRES2-Cre |
| 23 | human | 569818704 | 626170484 | MTG | 4 | NA |
| 24 | human | 569820026 | 626170475 | MTG | 5 | NA |

Detailed information about the aspiny/sparsely-spiny neuron models is provided below (the Index column corresponds to the left-to-right order in Figure 4. The neuron with index 10 is the sparsely-spiny neuron, and the rest are all aspiny.):

**Table 3:** Aspiny/sparsely-spiny model details.

| Index | Species | Cell ID | Model ID | Brain Region | Layer | Transgenic Line |
|---|---|---|---|---|---|---|
| 1 | mouse | 478793814 | 479694384 | VISp | 4 | Pvalb-IRES-Cre |
| 2 | mouse | 479220013 | 480056382 | VISp | 2/3 | Htr3a-Cre_NO152 |
| 3 | mouse | 488677994 | 489932551 | VISp | 2/3 | Sst-IRES-Cre |
| 4 | mouse | 484635029 | 485602029 | VISp | 2/3 | Pvalb-IRES-Cre |
| 5 | mouse | 484742372 | 485611914 | VISp | 5 | Pvalb-IRES-Cre |
| 6 | mouse | 488674910 | 489932183 | VISp | 6a | Ntsr1-Cre_GN220 |
| 7 | mouse | 475622680 | 478045202 | VISp | 2/3 | Htr3a-Cre_NO152 |
| 8 | human | 571709783 | 626170430 | MTG | 3 | NA |
| 9 | human | 531520637 | 626170592 | FroL | 3 | NA |
| 10 | human | 571718621 | 626170421 | MTG | 2 | NA |
| 11 | human | 529807751 | 626170661 | MTG | 3 | NA |

### B.3 Computational resources

All numerical experiments are conducted on a cluster equipped with two Intel(R) Xeon(R) Platinum 8582C CPUs and four NVIDIA GeForce RTX 5090 GPUs. The simulation of neurons is performed in parallel on this cluster, using CPUs, while all subsequent computations utilize only a single GPU.

## C Additional empirical summaries

### C.1 Aspiny/sparsely-spiny cells ranked by cell-level median SCI

In addition to the results of the spiny group reported in Figure 2**A**, we present here the results of the aspiny/sparsely-spiny group. SCI also separated human and mouse cells in the aspiny/sparsely-spiny group.

**Figure 4.**
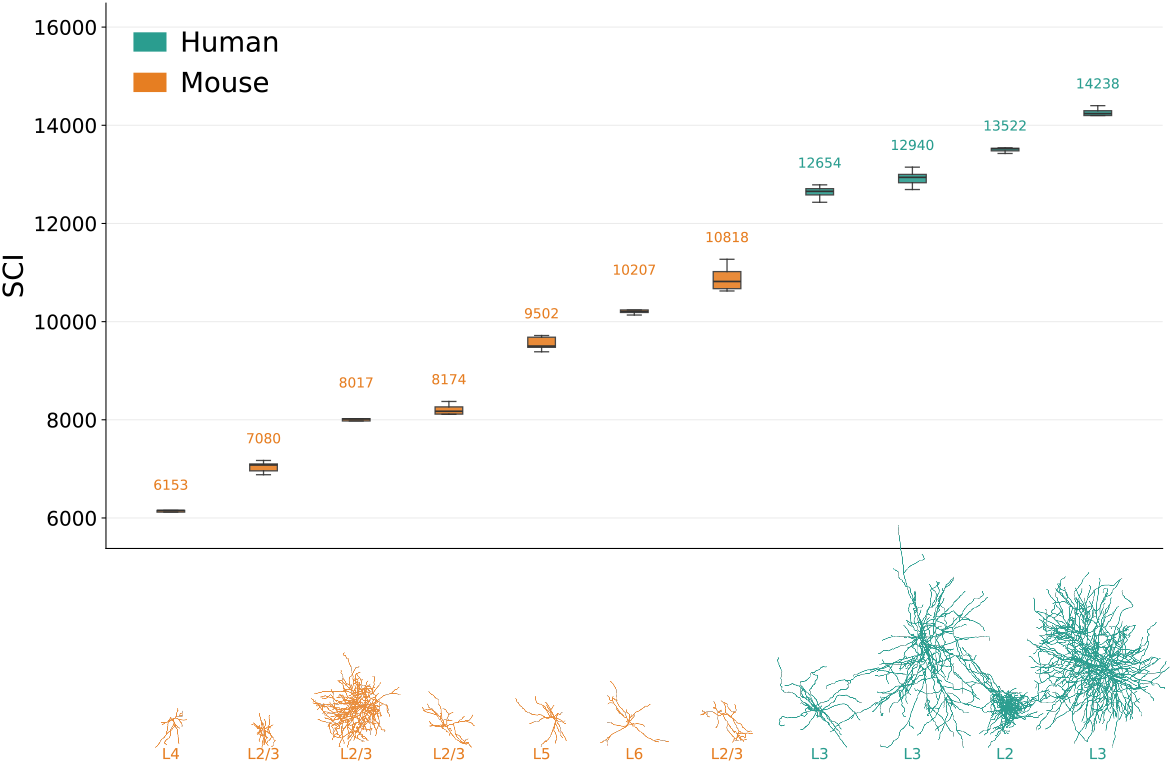
SCI separates human and mouse cortical aspiny/sparsely-spiny neuron models. Aspiny/sparsely-spiny cells are ranked by cell-level median 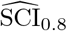. Boxplots summarize random-seed variability for each model (each box summarizes 5 SCI estimates from different window-sampling seeds), with morphology thumbnails and cortical-layer labels shown below. Human models are shown in teal and mouse models in orange. The second one from the right is a sparsely-spiny neuron, and the rest are all aspiny.

### C.2 Comparison of SCI between aspiny and spiny neurons

Biologically, the single-neuron computational complexity of aspiny neurons is overall lower than that of spiny neurons. SCI captures this phenomenon (K–S *D* = 0.610, *p* = 0.004; Welch *t* = −3.94, *p* = 0.002; Cohen’s *d* = 1.86; Cliff’s delta = 0.735).

**Figure 5.**
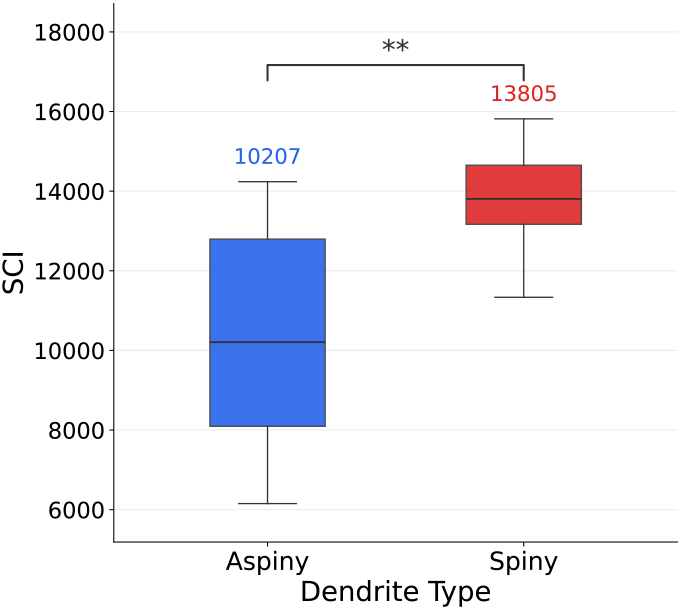
Spiny vs aspiny/sparsely-spiny comparison. Notice that the aspiny group contains the sparsely-spiny neuron. Statistical tests are performed across models rather than across random seeds, with each neuron using its median as a single sample point. Asterisks denote statistical significance according to the Kolmogorov–Smirnov (K–S) test (** *p <* 0.01).

### C.3 Subgroup agreement between SCI and FCI

The main text provides the SCI-FCI comparison for all cells in Figure 3, while here we present the SCI-FCI comparisons for four subsets. FCI and SCI maintain a statistically significant positive correlation across all four subsets.

**Figure 6.**
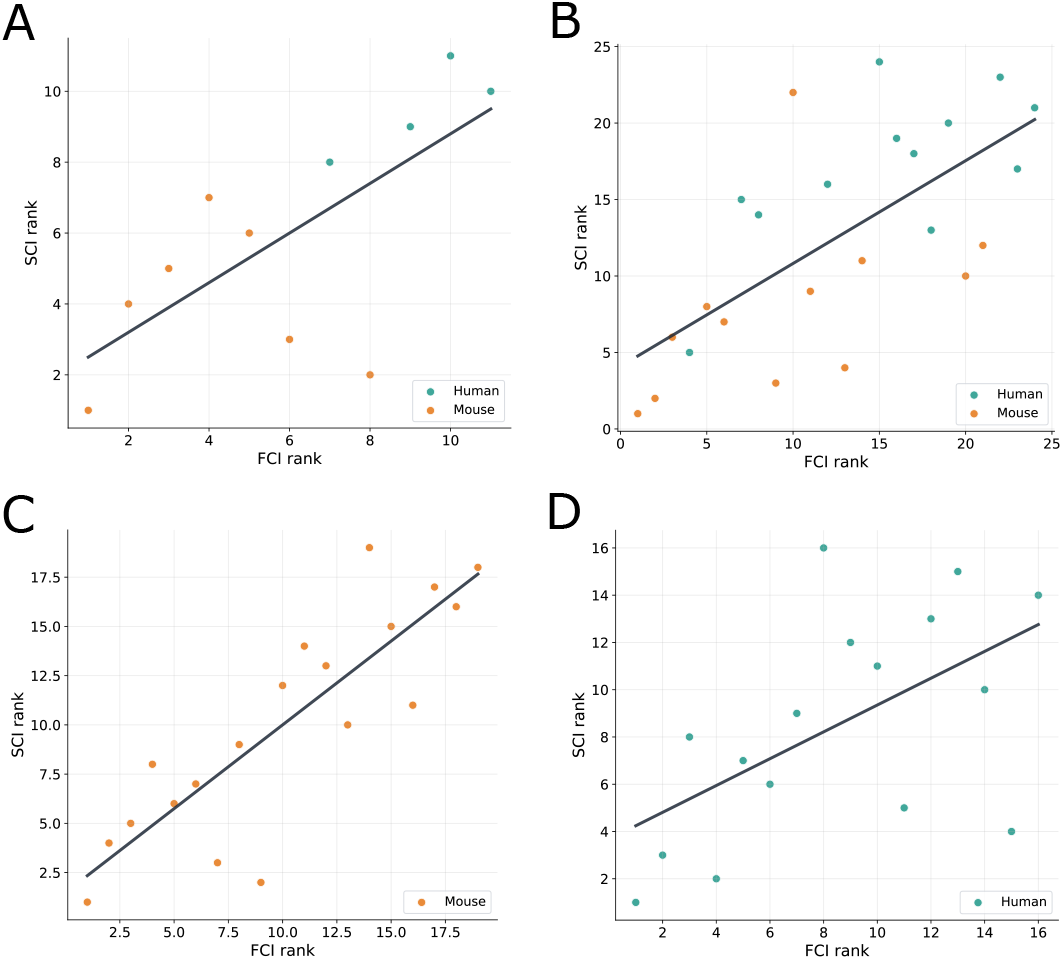
Subgroup agreement between SCI and FCI. The full-cell analysis in Figure 3 is decomposed into aspiny/sparsely-spiny, spiny, mouse, and human subsets. **A**, aspiny/sparsely-spiny subset. **B**, spiny subset. **C**, mouse subset. **D**, human subset. Spearman correlations are positive in all subsets: aspiny/sparsely-spiny *ρ* = 0.700, spiny *ρ* = 0.672, mouse *ρ* = 0.851, and human *ρ* = 0.568.

### C.4 Changes in SCI following NMDA receptor blockade

SCI captures the expected reduction in computational complexity following NMDA receptor blockade. The NMDA OFF condition used the same simulation pipeline as the main analysis, with the NMDA conductance ratio set to zero. The reduction in SCI is statistically significant under one-sided Wilcoxon signed-rank tests.

For mean SCI, Wilcoxon statistic *V* = 1, one-sided *p* = 5 ×10^*−*4^, Cohen’s *d* = −1.19; for median SCI, Wilcoxon statistic *V* = 2, one-sided *p* = 7 ×10^*−*4^, Cohen’s *d* = −1.25. This result provides additional validation for using SCI as a metric for single-neuron computational complexity, as NMDA receptors are known to be involved in nonlinear dendritic integration and complex input–output transformations. Notably, even after the removal of NMDA receptors, human cortical pyramidal neurons remain more complex than mouse cortical pyramidal neurons with intact NMDA receptors; this indicates that human cortical pyramidal neurons still possess a multitude of other specific mechanisms to maintain nonlinearity and complexity.

**Figure 7.**
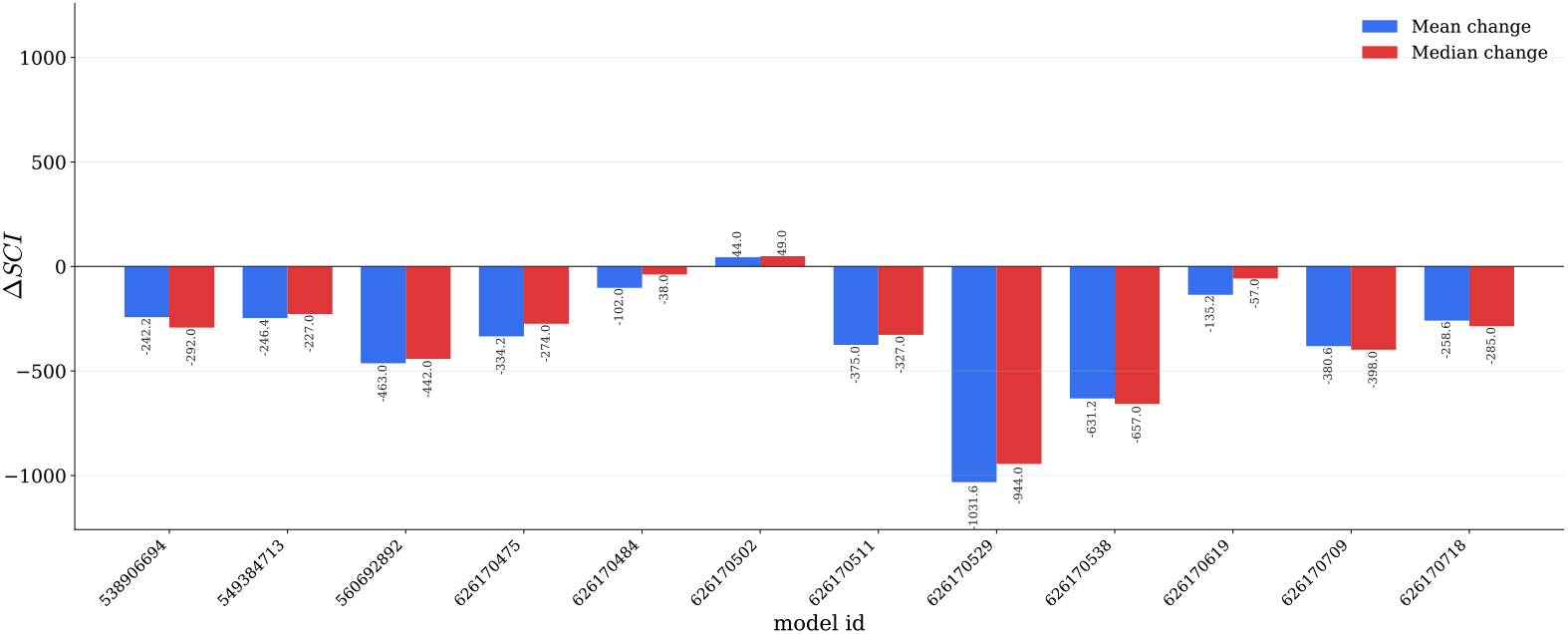
SCI captures the expected reduction in computational complexity following NMDA receptor blockade. We conducted experiments on all 12 human spiny models, with the x-axis representing the Model ID; see Table 2 for detailed information.

### C.5 Associations between SCI and morphology

**Figure 8.**
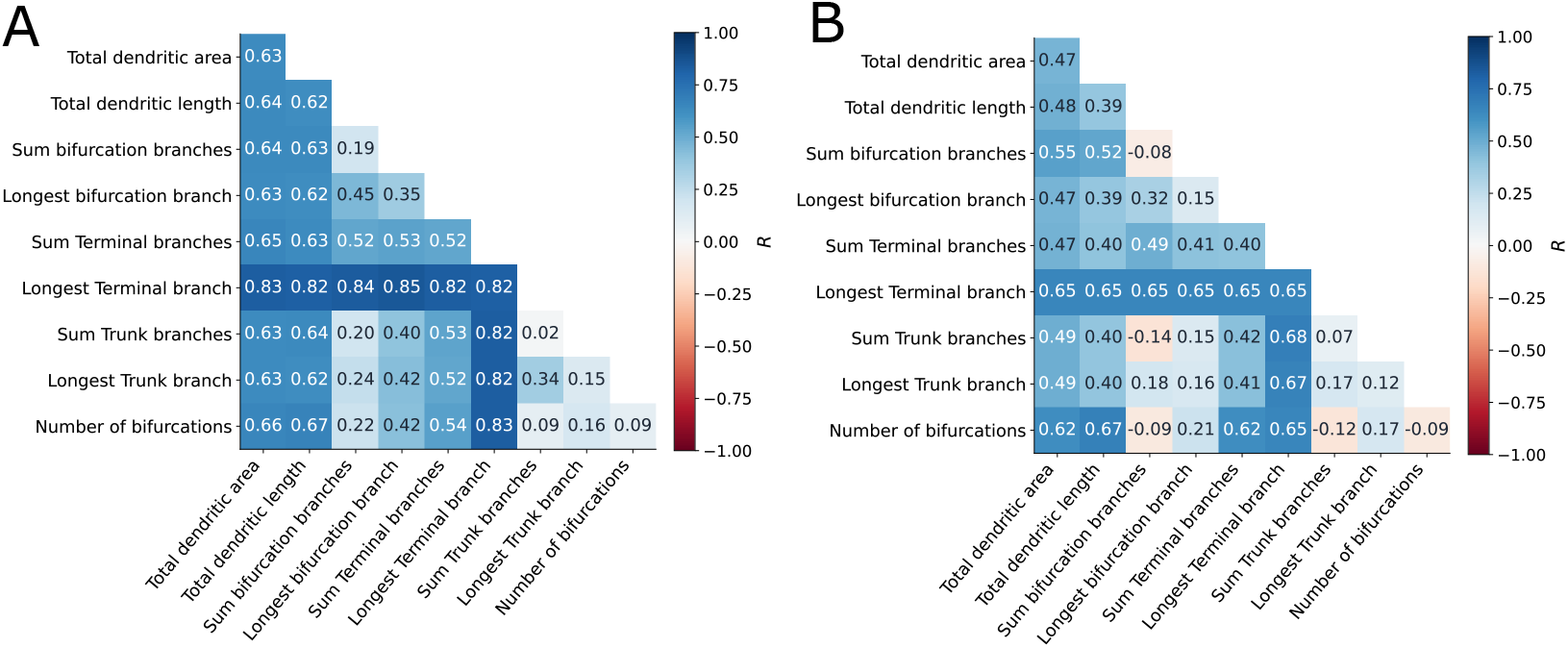
Associations between SCI and basal dendritic morphology summaries. **A–B**, Pairwise rank-regression analysis relating model-level median 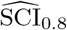 to dendritic morphology features in aspiny/sparsely-spiny and spiny cells, respectively. Diagonal entries show the *R* obtained from a single morphology feature, and off-diagonal entries show the *R* obtained from the corresponding pair of features. Darker blue indicates larger positive *R*; red indicates negative *R*. These analyses summarize associations with measured gross morphology features and are not used as causal evidence for the origin of SCI differences.

### C.6 Performance of the linear baseline

To demonstrate the necessity of using a weighted Gaussian kernel, we evaluated a linear baseline 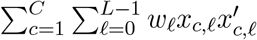. All other settings are identical to the default settings in the main text. The results indicated that the linear kernel completely failed to distinguish between human and mouse cortical neuron models. For the aspiny/sparsely-spiny group, K–S *D* = 0.5, *p* = 0.42, Welch *t* = 0.62, *p* = 0.57; for the spiny group, K–S *D* = 0.25, *p* = 0.87, Welch *t* = −1.23, *p* = 0.23.

**Figure 9.**
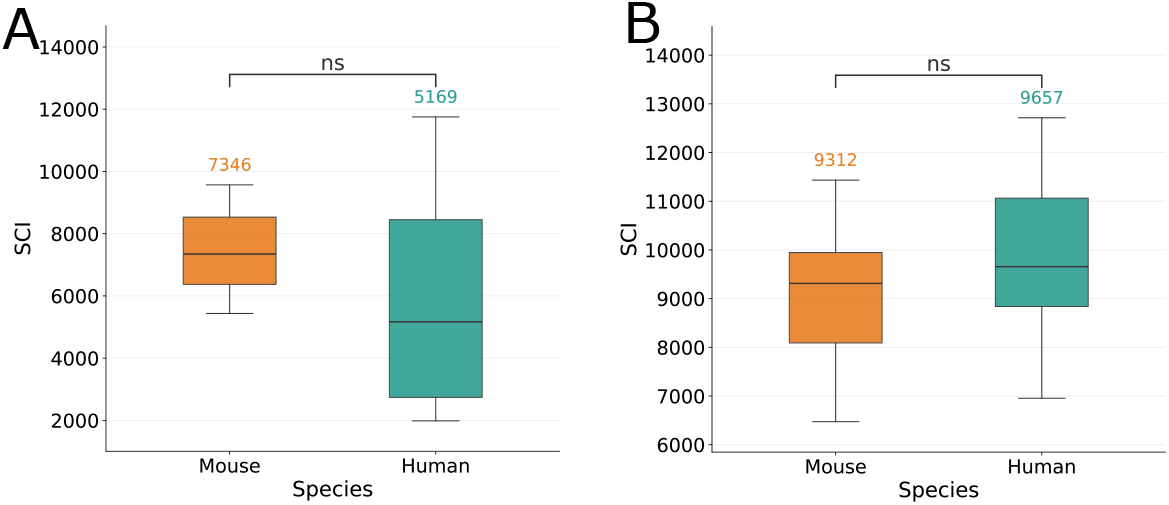
**A–B**, Species-level summaries using model medians for aspiny/sparsely-spiny cells and spiny cells, respectively. Statistical tests are performed across models rather than across random seeds, with each neuron using its median as a single sample point. ns denotes not statistically significant according to the Kolmogorov–Smirnov (K–S) test (ns, *p >* 0.05).

### C.7 SCI robustness

**Table 4:** Rank robustness of SCI relative to the baseline setting (*σ* = 1, *τ* = 20 ms, *L* = 200 ms, Δ_stride_ = 50 ms, *θ* = 0.8, *M* = 16384). Spearman correlations are computed over cell-level SCI ranks.

| Perturbation | Spearman *ρ* | *p*-value |
|---|---|---|
| *M* = 4096 | 0.971 | 4.7 × 10<sup>−22</sup> |
| *M* = 8192 | 0.989 | 3.3 × 10<sup>−29</sup> |
| *M* = 12288 | 0.994 | 4.1 × 10<sup>−33</sup> |
| Use only 100 10-second simulations | 0.992 | 1.2 × 10<sup>−30</sup> |
| Use only 10 10-second simulations, Δ<sub>stride</sub> = 5 ms | 0.974 | 3.9 × 10<sup>−22</sup> |
| *L* = 50 ms, Δ<sub>stride</sub> = 50 ms | 0.994 | 3.3 × 10<sup>−32</sup> |
| *L* = 100 ms, Δ<sub>stride</sub> = 100 ms | 0.993 | 1.5 × 10<sup>−31</sup> |
| *L* = 500 ms, Δ<sub>stride</sub> = 200 ms | 0.997 | 5.2 × 10<sup>−37</sup> |
| Centered kernel | 0.995 | 3.4 × 10<sup>−34</sup> |
| Unnormalized electrotonic distance | 0.972 | 9.6 × 10<sup>−22</sup> |
| *σ* = 0.5 | −0.191 | 0.28 |
| *σ* = 1.5 | 0.980 | 4.7 × 10<sup>−24</sup> |
| *σ* = 2.0 | 0.934 | 8.0 × 10<sup>−16</sup> |
| *τ* = 10 ms | 0.991 | 3.6 × 10<sup>−29</sup> |
| *τ* = 30 ms | 0.987 | 3.1 × 10<sup>−27</sup> |
| *τ* = 50 ms | 0.367 | 0.033 |
| *θ* = 0.6 | 0.789 | 3.0 × 10<sup>−8</sup> |
| *θ* = 0.65 | 0.900 | 4.6 × 10<sup>−13</sup> |
| *θ* = 0.7 | 0.982 | 1.3 × 10<sup>−24</sup> |
| *θ* = 0.75 | 0.997 | 5.2 × 10<sup>−37</sup> |
| *θ* = 0.85 | 0.990 | 6.0 × 10<sup>−29</sup> |
| *θ* = 0.9 | 0.973 | 5.7 × 10<sup>−22</sup> |

## References

[1] Panayiota Poirazi, Terrence Brannon, and Bartlett W Mel. “Pyramidal neuron as two-layer neural network”. In: Neuron 37.6 (2003), pp. 989–999. DOI:10.1016/S0896-6273(03)00149-1.

[2] Alon Polsky, Bartlett W Mel, and Jackie Schiller. “Computational subunits in thin dendrites of pyramidal cells”. In: Nature neuroscience 7.6 (2004), pp. 621–627. DOI:10.1038/nn1253.

[3] Tiago Branco, Beverley A Clark, and Michael Häusser. “Dendritic discrimination of temporal input sequences in cortical neurons”. In: Science 329.5999 (2010), pp. 1671–1675. DOI:10.1126/science.1189664.

[4] Michael London and Michael Häusser. “DENDRITIC COMPUTATION”. In: Annual Review of Neuroscience 28 (2005), pp. 503–532. ISSN: 1545-4126. DOI:10.1146/annurev.neuro.28.061604.135703.

[5] Guy Major, Matthew E. Larkum, and Jackie Schiller. “Active Properties of Neocortical Pyramidal Neuron Dendrites”. In: Annual Review of Neuroscience 36 (2013), pp. 1–24. ISSN: 1545-4126. DOI:10.1146/annurev-neuro-062111-150343.

[6] Hemanth Mohan, Matthijs B Verhoog, Keerthi K Doreswamy, Guy Eyal, Romy Aardse, Brendan N Lodder, Natalia A Goriounova, Boateng Asamoah, AB Clementine B. Brakspear, Colin Groot, et al. “Dendritic and axonal architecture of individual pyramidal neurons across layers of adult human neocortex”. In: Cerebral cortex 25.12 (2015), pp. 4839–4853. DOI:10.1093/cercor/bhv188.

[7] Guy Eyal, Matthijs B Verhoog, Guilherme Testa-Silva, Yair Deitcher, Johannes C Lodder, Ruth Benavides-Piccione, Juan Morales, Javier DeFelipe, Christiaan PJ de Kock, Huibert D Mansvelder, et al. “Unique membrane properties and enhanced signal processing in human neocortical neurons”. In: elife 5 (2016), e16553. DOI:10.7554/eLife.16553.

[8] Guy Eyal, Matthijs B. Verhoog, Guilherme Testa-Silva, Yair Deitcher, Ruth Benavides-Piccione, Javier DeFelipe, Christiaan P. J. de Kock, Huibert D. Mansvelder, and Idan Segev. “Human Cortical Pyramidal Neurons: From Spines to Spikes via Models”. In: Frontiers in Cellular Neuroscience Volume 12-2018 (2018). ISSN: 1662-5102. DOI: 10.3389/fncel.2018.00181. URL: https://www.frontiersin.org/journals/cellular-neuroscience/articles/10.3389/fncel.2018.00181.

[9] Albert Gidon, Timothy Adam Zolnik, Pawel Fidzinski, Felix Bolduan, Athanasia Papoutsi, Panayiota Poirazi, Martin Holtkamp, Imre Vida, and Matthew Evan Larkum. “Dendritic action potentials and computation in human layer 2/3 cortical neurons”. In: Science 367.6473 (2020), pp. 83–87. DOI: 10.1126/science.aax6239.

[10] David Beniaguev, Idan Segev, and Michael London. “Single cortical neurons as deep artificial neural networks”. In: Neuron 109.17 (2021), pp. 2727–2739. DOI:10.1016/j.neuron.2021.07.002.

[11] Ido Aizenbud, Daniela Yoeli, David Beniaguev, Christiaan PJ de Kock, Michael London, and Idan Segev. “What makes human cortical pyramidal neurons functionally complex”. In: bioRxiv (2024). DOI:10.1101/2024.12.17.628883.

[12] Jingyang Ma, Songting Li, and Douglas Zhou. “Mapping Biological Neuron Dynamics into an Interpretable Two-layer Artificial Neural Network”. In: arXiv preprint arXiv:2305.12471 (2023). DOI: 10.48550/arXiv.2305.12471.

[13] Oren Amsalem, Guy Eyal, Noa Rogozinski, Michael Gevaert, Pramod Kumbhar, Felix Schürmann, and Idan Segev. “An efficient analytical reduction of detailed nonlinear neuron models”. In: Nature communications 11.1 (2020), p. 288. DOI:10.1038/s41467-019-13932-6.

[14] Willem AM Wybo, Jakob Jordan, Benjamin Ellenberger, Ulisses Marti Mengual, Thomas Nevian, and Walter Senn. “Data-driven reduction of dendritic morphologies with preserved dendro-somatic responses”. In: Elife 10 (2021), e60936. DOI:10.7554/eLife.60936.

[15] Bernhard Schölkopf, Alexander Smola, and Klaus-Robert Müller. “Nonlinear component analysis as a kernel eigenvalue problem”. In: Neural computation 10.5 (1998), pp. 1299–1319. DOI:10.1162/089976698300017467.

[16] Ingo Steinwart and Clint Scovel. “Mercer’s theorem on general domains: On the interaction between measures, kernels, and RKHSs”. In: Constructive Approximation 35.3 (2012), pp. 363–417. DOI:10.1007/s00365-012-9153-3.

[17] Michael L Hines and Nicholas T Carnevale. “The NEURON simulation environment”. In: Neural computation 9.6 (1997), pp. 1179–1209. DOI:10.1162/neco.1997.9.6.1179.

[18] Nicholas T Carnevale and Michael L Hines. The NEURON book. Cambridge University Press, 2006.

[19] Allen Institute for Brain Science. Allen Cell Types Database – Human Morphology-Electrophysiology. Dataset. RRID:SCR_0148062015. URL: https://celltypes.brain-map.org/data.

[20] Nathan W Gouwens, Staci A Sorensen, Jim Berg, Changkyu Lee, Tim Jarsky, Jonathan Ting, Susan M Sunkin, David Feng, Costas A Anastassiou, Eliza Barkan, et al. “Classification of electrophysiological and morphological neuron types in the mouse visual cortex”. In: Nature neuroscience 22.7 (2019), pp. 1182–1195. DOI:10.1038/s41593-019-0417-0.

[21] Lida Kanari, Paweł Dłotko, Martina Scolamiero, Ran Levi, Julian Shillcock, Kathryn Hess, and Henry Markram. “A topological representation of branching neuronal morphologies”. In: Neuroinformatics 16.1 (2018), pp. 3–13. DOI:10.1007/s12021-017-9341-1.

[22] Bojan Mihaljević, Pedro Larrañaga, and Concha Bielza. “Comparing the electrophysiology and morphology of human and mouse layer 2/3 pyramidal neurons with bayesian networks”. In: Frontiers in neuroinformatics 15 (2021), p. 580873. DOI:10.3389/fninf.2021.580873.

[23] Anna A Galakhova, Sarah Hunt, René Wilbers Djai B Heyer, Christiaan PJ de Kock, Huibert D Mansvelder, and Natalia A Goriounova. “Evolution of cortical neurons supporting human cognition”. In: Trends in cognitive sciences 26.11 (2022), pp. 909–922. DOI:10.1016/j.tics.2022.08.012.

[24] Sahil Loomba, Jakob Straehle, Vijayan Gangadharan, Natalie Heike, Abdelrahman Khalifa, Alessandro Motta, Niansheng Ju, Meike Sievers, Jens Gempt, Hanno S Meyer, et al. “Connectomic comparison of mouse and human cortex”. In: Science 377.6602 (2022), eabo0924. DOI:10.1126/science.abo0924.

[25] Homeira Moradi Chameh, Madeleine Falby, Mandana Movahed, Keon Arbabi, Scott Rich, Liang Zhang, Jérémie Lefebvre, Shreejoy J Tripathy, Maurizio De Pittà, and Taufik A Valiante. “Distinctive biophysical features of human cell-types: insights from studies of neurosurgically resected brain tissue”. In: Frontiers in synaptic neuroscience 15 (2023), p. 1250834. DOI: 10.3389/fnsyn.2023.1250834.

[26] Spyridon Chavlis and Panayiota Poirazi. “Dendrites endow artificial neural networks with accurate, robust and parameter-efficient learning”. In: Nature communications 16.1 (2025), p. 943. DOI:10.1038/s41467-025-56297-9.

[27] James Mercer. “Xvi. functions of positive and negative type, and their connection the theory of integral equations”. In: Philosophical transactions of the royal society of London. Series A, containing papers of a mathematical or physical character 209 (1909), pp. 415–446. DOI:10.1098/rsta.1909.0016.

